# The *LYCOPENE EPSILON CYCLASE* untranslated mRNA leader modulates carotenoid feedback and post-transcriptional regulation

**DOI:** 10.1101/2024.07.19.604344

**Authors:** Yagiz Alagoz, Jwalit J. Nayak, Rishi Aryal, Jacinta L. Watkins, Sophie Holland, David T. Tissue, Barry J. Pogson, Christopher I. Cazzonelli

## Abstract

Metabolic feedback is proposed to modulate nuclear gene expression and carotenoid biosynthesis in plastids, however few mechanisms have been identified so far in plants. Utilising mutants, overexpression lines, and chemical inhibitors, we demonstrate that Arabidopsis *LYCOPENE EPSILON CYCLASE* (*εLCY*) mRNA levels correlate with changes in β-carotenoid accumulation. Transgenic seedlings harbouring the *εLCY* 5’ leader sequence fused to *FIREFLY LUCIFERASE* (*FiLUC*) showed reporter responsiveness to metabolic feedback triggered by norflurazon or loss-of-function in the CAROTENOID ISOMERASE (CRTISO). The *εLCY* 5’UTR harboured three alternative transcription start sites (TSS). The most abundant -133bp sequence generated in dark and light grown seedlings harboured a 5’ conserved domain (CD) with other *Brassicaceae* species and a viral internal ribosome entry site (IRES) proximal to the start codon. *In silico* modelling predicted the 5’UTR formed two energetically separated RNA structural probabilities having a minimal free energy consistent with metabolite-binding RNA riboswitches that was distinguished by hairpin structures within the CD. Site-specific mutations were used to stabilize the 5’UTR into a single RNA shape definition having negligible separation between the mountain plot structure prediction curves and a distal terminator-like hairpin structure. Stabilizing the 5’UTR shape triggered the posttranscriptional repression of FiLUC activity enabled by the CaMV35S promoter in tobacco transient assays and stable transgenic Arabidopsis lines. The stabilised shape fragment became responsive to metabolic feedback induced by norflurazon and in *crtiso* mutant etiolated and de-etiolated seedlings. The *εLCY* 5’UTR resembles a conformational RNA regulatory switch harbouring a posttranscriptional expression platform and aptamer domain responsive to carotenoid-mediated feedback signalling.

## INTRODUCTION

Carotenoids are secondary metabolites that serve as pigments contributing to photosynthesis, photoprotection, and are substrates for the generation of apocarotenoid signalling metabolites, as well as phytohormones (Baranski and Cazzonelli, 2016; Sun et al., 2022). Carotenoid biosynthesis, accumulation, and catabolism in plastids and tissues are tightly controlled by metabolic feedback mechanisms that arise from fluctuations in carotenoid levels due to genetic perturbations, environmental change (e.g., light, temperature, and water stress), and developmental transitions (Cazzonelli and Pogson, 2010; Llorente et al., 2017; Sun et al., 2018). For example, the promoter and/or 5’UTR can coordinate the transcriptional and posttranscriptional processes, yet these regulatory processes have not been well established for most carotenoid genes. In Arabidopsis, the 5’UTR of *PHYTOENE SYNTHASE* (*PSY*) mRNA was shown to control carotenogenesis by altering the abundance of *PSY* splice variants during different light conditions and displayed RNA structural characteristics reminiscent of an RNA regulatory switch (Álvarez et al., 2016). Yet, mechanisms for such transcriptional changes across the pathway remain largely elusive.

Carotenoids are precursors of some phytohormones such as abscisic acid (ABA) and strigolactones (SL) as well as various apocarotenoids (Moreno et al., 2021; Zheng et al., 2021). Transcriptional responses associated with any carotenoid cleavage product, herein referred to as an apocarotenoid signal (ACS) have been linked to the control of carotenoid metabolic flux, plastid biogenesis and differentiation leaf and root development, photo-acclimation, plant root-mycorrhizal interactions, herbivore defence, and plant growth (Van Norman et al., 2014; Dickinson et al., 2019; Cazzonelli et al., 2020; Escobar-Tovar et al., 2021; Nayak et al., 2022; Hou et al., 2024). A linear *cis*-carotene-derived apocarotenoid signal (*cis*-ACS) was shown to regulate key repressor (e.g., PHYTOCHROME INTERACTING FACTOR3; PIF3) and activator (e.g., ELONGATED HYPOCOTYL5; HY5) transcription factors involved in controlling *PHOTOSYNTHESIS ASSOCIATED NUCLEAR GENE* (*PhANG*) expression during the dark to light transition (Cazzonelli et al., 2020). Establishing the physiological and/or biochemical basis to a threshold level of the carotenoid substrate(s) that regulate metabolic feedback responses can be a challenge since the substrate(s) are more than quantities normally required to produce the bioactive signal (Hou et al., 2024).

Metabolic feedback could involve an interaction between a mobile bioactive metabolic signal and ligand-binding sensory domain (messenger RNA, DNA promoter *cis*-acting element, and/or transcription factor/enzyme active site) that causes aptamer alterations in structure, DNA-protein binding, or protein-protein interactions, leading to changes in transcriptional or translational expression platforms that modulate metabolite biosynthesis (Alagoz et al., 2018; Agrawal et al., 2022; Kavita and Breaker, 2022). Molecular models have proposed that an ACS could block enzyme catalysis, alter intron splicing, or affect protein-protein interactions (Sun et al., 2014; Álvarez et al., 2016; Mitra et al., 2021). The targets of metabolic feedback and how a plant-derived ACS regulates transcription or posttranscriptional processes remain to be elucidated.

LYCOPENE EPSILON CYCLASE (εLCY) regulates the α-branch of the carotenoid biosynthetic pathway and is a rate-limiting step in the production of lutein, the most abundant carotenoid in foliar tissues (Cunningham Jr et al., 1996; Pogson et al., 1996). The downregulation of *εLCY* gene expression by genetic modifications or loss-of-function of εLCY activity redirects flux towards the β-branch in the pathway (10 out 14 published reports; Table S1). This has been adopted as a biofortification strategy to enhance β-carotene (pro-vitamin A) accumulation in crops (Diretto et al., 2006; Yu et al., 2008; Kim et al., 2013; Shi et al., 2014; Ke et al., 2019; Kaur et al., 2020).

Intriguingly, modulation of precursor synthesis through the alpha- and beta-branches of the pathway changes flux and *εLCY* transcript levels (Table S1). That is, exogenous expression of the *Pantoea ananatis* phytoene desaturase (CRTI) gene in wild-type tomato, *tangerine,* and *old gold crimson* enhanced β-carotenoids and *εLCY* expression during fruit ripening (Enfissi et al., 2017) (Table S1). Similarly, dark-grown Arabidopsis *ccr2* (*carotenoid* c*hloroplast regulator 2*) seedlings that lack the function of the CAROTENOID ISOMERASE (CRTISO) do not biosynthesize cyclic carotenoids, accumulate linar *cis*-carotenes, and exhibit a negative regulation in *εLCY* expression (Park et al., 2002; Cuttriss et al., 2007). *εLCY* expression was also reduced in older mature rosette leaves from the *ccr1* (*carotenoid* c*hloroplast regulator 1*) mutant that lacks function in SET DOMAIN GROUP 8 (SDG8) that is required to maintain *CRTISO* expression (Cazzonelli et al., 2009). The genetic, environmental, and developmental changes that trigger feedback regulation of *εLCY* expression, and how they correlate with fluctuations in carotenogenesis (e.g., acyclic *cis*-carotenes vs cyclic carotenoids) remain to be discovered. The molecular mechanisms modulating *εLCY* gene expression in the nucleus have yet to be defined.

The UTRs of some prokaryote genes can harbour RNA structural domains that sense/bind metabolites (e.g., metabolite-binding RNA riboswitches), change conformational shape in response to temperature shifts (e.g., RNA thermoswitches), and modulate translation initiation (e.g. internal ribosome entry site; IRES) of which one is conserved in eukaryotes (e.g. *THIAMINE PYROPHOSPHATE; TPP* riboswitch) (Altuvia et al., 1989; Dinkova et al., 2005; Bocobza et al., 2007; Wachter et al., 2007; Li and Breaker, 2013; Jiménez-González et al., 2014; Cui et al., 2015; Chung et al., 2020). Changes in the transcription start site (TSS) modify the length of 5’UTR, splicing, presence/absence of regulatory motifs, and RNA structural formation that impacts gene transcription, mRNA decay, and/or protein translation (Thatcher et al., 2007; Álvarez et al., 2016). Transposon insertions and allelic variation in the 5’ upstream regions adjacent to the *εLCY* mRNA start codon can determine flux between the lutein and β-carotene branches, leading to β-carotene accumulation in maize and wheat (Harjes et al., 2008; Muthusamy et al., 2015; Richaud et al., 2018).

Here, we undertook a genetic, environmental (light), chemical (norflurazon; NFZ) and developmental (etiolated vs de-etiolated seedlings, and emerging leaves) approach to decipher the signals that regulate *εLCY* expression *in planta* and determine if they correlate with feedback regulation, due to the accumulation of any specific carotenoid substrate. We engineered promoter-reporter gene fusions and utilised stable transgenics as well as transient expression systems to investigate the mechanism by which metabolic feedback signals can regulate an exogenously expressed *εLCY* promoter-luciferase transgene together with endogenous *εLCY* mRNA transcription in tissues enriched in etioplasts (dark) or chloroplasts (light). We hypothesised that the *εLCY* mRNA 5’ leading sequence contributes to the feedback regulation of *εLCY* transcription and searched for alternative transcription initiation sites and conserved sequence domains among *Brassicaceae* species. Bioinformatics modelling of RNA structural definitions combined with mutational/gain-of-function strategies were used to interrogate the impact of *εLCY* 5’UTR shape, regulatory motifs, and alternative TSS on modulating *εLCY* promoter-luciferase mRNA and reporter activity levels *in planta* towards gaining insights into whether the *εLCY* 5’UTR could function like an RNA regulatory switch in response to metabolic feedback.

## RESULTS

### Accumulation of linear *cis*-carotenes does not regulate *εLCY* expression

We utilised carotenoid biosynthetic mutants accumulating linear *cis*-carotenes in dark-grown seedlings to decipher if they trigger feedback regulation of *εLCY* expression. The tomato Micro-Tom *tangerine* (*tang^Mic^*) and *Arabidopsis ccr2* mutants that lack CRTISO activity and accumulate various *cis*-carotenes (Isaacson et al., 2002; Park et al., 2002) show reduced *εLCY* transcript levels in dark grown seedlings (Figure 1A-B). The loss-in-function of ZETA-CAROTENE ISOMERASE (*ziso*) in the *Arabidopsis ccr2 ziso* double mutant prevented the synthesis of neurosporene and tetra-*cis*-lycopene, restored prolamellar body (PLB) formation in *ccr2* (Cazzonelli et al., 2020), and reduced total *cis*-carotene levels by 50% (Figure S1A). *ccr2*, *ziso*, and *ccr2 ziso* all show a similar 2-fold reduction in *εLCY* expression (Figure 1B). The loss-in-function of DE-ETIOLATED 1 (DET1) in *ccr2 det1-154* dark grown seedlings reduced di-*cis*-ζ-carotene, neurosporene, and tetra-*cis*-lycopene levels as well as restored PLB formation in *ccr2* (Cazzonelli et al., 2020). *ccr2 det1-154* etiolated seedlings exhibit a similar 80% reduction in *εLCY* expression when compared to *det1-1* (Figure 1C). Therefore, neither the accumulation of *cis*-carotenes or their derivative signalling functions over PLB formation in *ccr2* appear to regulate *εLCY* expression.

**Figure 1.**
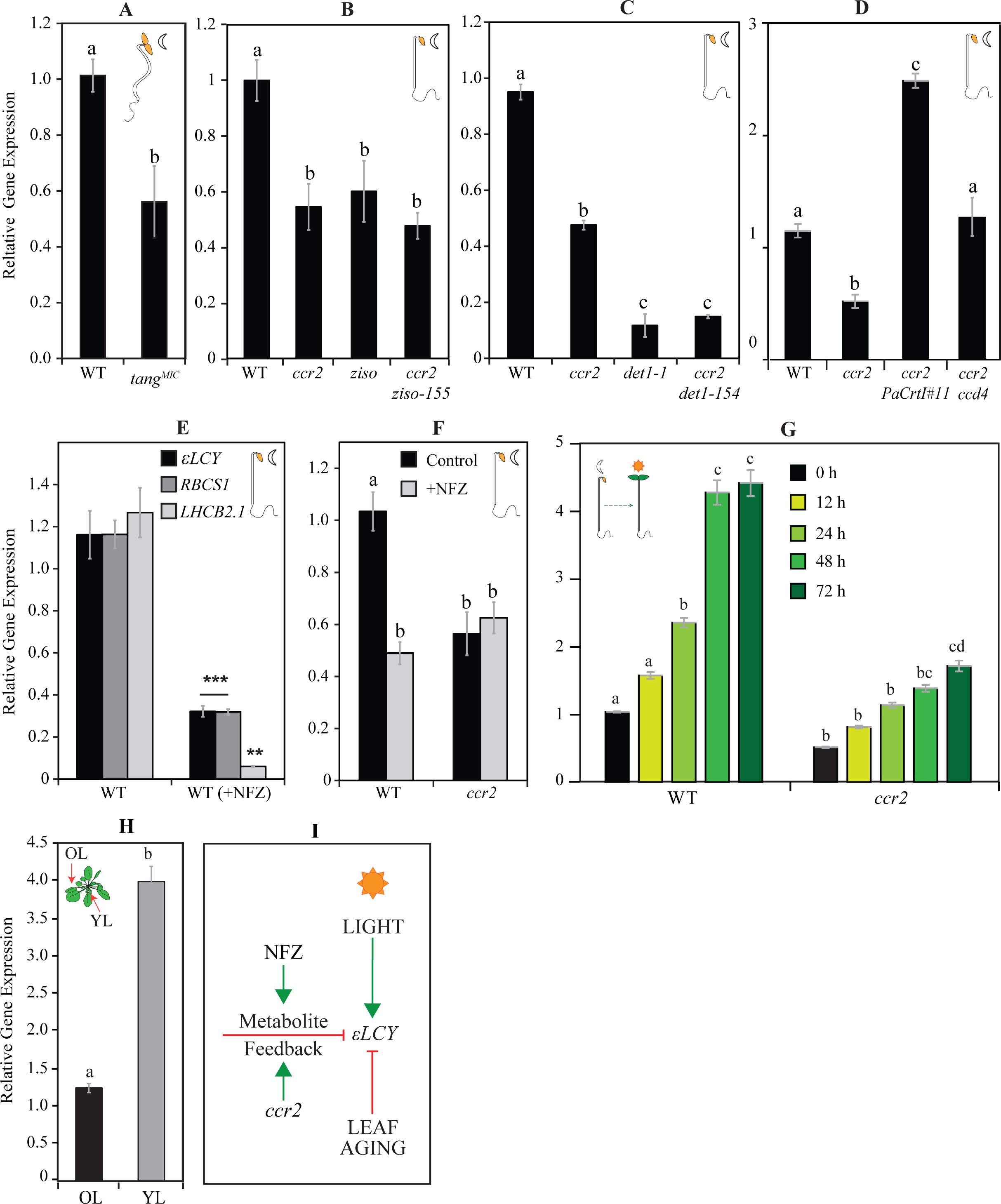
Regulation of *εLCY* expression by chemical, genetic, developmental, and environmental changes in seedling and leaf tissues. **A-C)** *εLCY* expression in linear *cis*-carotene accumulating etiolated tissues from (**A**) MicroTom *tangerine* (*tang^Mic^*), (**B**) *ccr2*, *ziso*, and *ccr2 ziso*, and (**C**) *det1-154*, *ccr2*, and *ccr2 det1-154*. **D**) *εLCY* expression in WT, *ccr2*, *ccr2 T35ehn*::*PaCrtI#11* (*ccr2* transformed with pT35enh::SSU-*PaCrtI*), and *ccr2 ccd4* etiolated seedlings that accumulate linear *cis*-carotenes plus cyclic carotenoids. **E**) Relative expression of photosynthesis-associated nuclear genes (*RBCS1*, *LHCB2.1*) and *εLCY* in WT etiolated tissues treated with NFZ. **F**) *εLCY* expression in *ccr2* etiolated tissues treated with NFZ. **G**) *εLCY* expression in WT and *ccr2* etiolated and de-etiolated seedlings exposed to continuous light for up to 72 hrs. **H**) *εLCY* expression in young emerging leaves (YL) and older mature leaves (OL) from whole rosettes. **I**) Model summarising feedback regulations of *εLCY* expression triggered by chemical (NFZ), genetic (*ccr2*), developmental (leaf age) and environmental (light) perturbations. Data is representative of two to three independent experiments, and standard error bars of the mean are displayed (n=3-9). Lettering denotes significance by one- or two-way ANOVA statistical analysis with post-hoc Tukey test. Abbreviations: *εLCY*; *LYCOPENE EPSILON CYCLASE*, *RBCS1; RIBULOSE BISPHOSPHATE CARBOXYLASE SMALL CHAIN 1A, LHCB2.1; LIGHT-HARVESTING CHLOROPHYLL B-BINDING 2*

### Changes in cyclic carotenoids can feedback to regulate *εLCY* expression

We next queried if changes in cyclic carotenoids downstream of linear *cis*-carotene biosynthesis can feedback to regulate *εLCY* expression. *εLCY* expression was reduced by ∼80% in *det1-1* etiolated seedlings (Figure 1C) previously shown to accumulate 3-fold less cyclic carotenoids compared to WT (Cazzonelli et al., 2020). The biosynthesis of all-*trans*-lycopene can bypass *cis*-carotene precursors in plants by exogenously expressing *Pantoea agglomerans* (*Erwinia herbicola*) or *Pantoea ananatis* (*Erwinia uredovora*) *CAROTENOID ISOMERASE* (*PaCrtI*) (Romer et al., 2000; Yoon et al., 2007; Enfissi et al., 2017). A transgene harbouring *PaCrtI* under control by the CaMV35s promoter (*T35enh*::*SSU-PaCrtI*) was transformed into *ccr2* to enhance all-*trans*-lycopene biosynthesis in etiolated *ccr2* seedlings. Four homozygous *ccr2 35S::PaCrtI* lines (#5, 6, 9, and 11) that segregated in a typical 3:1 manner (Chi^2 *p*-value > 0.64) displayed a carotenoid composition similar to that of *ccr2* when grown under a long 16 h photoperiod (Table S2). Etiolated tissues from *ccr2 T35enh::SSU-PaCrtI*#11 showed a 2- to 3-fold increase in neoxanthin, violaxanthin, antheraxanthin, and β-carotene as well as 2.3- to 3.8-fold reduction in linear *cis*-carotene levels (Figure S1B-C). *ccr2 T35enh::SSU-PaCrtI*#11 etiolated seedlings displayed a 5-fold and 2.4-fold higher *εLCY* transcript abundance compared to *ccr2* and WT, respectively (Figure 1D). Therefore, the increased cyclic carotenoid levels in *PaCrtI* exogenous expression in etiolated *ccr2* tissues can enhance *εLCY* transcript levels.

CAROTENOID CLEAVAGE DIOXYGENASE 4 (CCD4) modulates β-carotene accumulation in Arabidopsis as *ccd4* mutants show enhanced β-carotene accumulation in seeds and senescing leaves (Næsted et al., 2004; Gonzalez-Jorge et al., 2013; Bhuiyan et al., 2016). We tested if *ccd4* introgressed with *ccr2* would enrich cyclic carotenoid levels and alter *εLCY* mRNA levels in etiolated tissues. Compared to *ccr2*, the *ccr2 ccd4* etiolated seedlings had a similar linear *cis*-carotene abundance yet significantly higher levels of β-carotene and zeaxanthin (Figure S2A-B). *ccr2 ccd4* etiolated tissues showed a 2-fold increase in *εLCY* expression relative to *ccr2* (Figure 1D). Taken together, reductions and increases in cyclic carotenoids (rather than linear *cis*-carotenes) in etiolated seedlings trigger negative and positive feedback regulation of *εLCY* expression, respectively.

### Chemical inhibition of carotenoid biosynthesis suppresses *εLCY* expression

Norflurazon (NFZ) chemically inhibits PDS enzyme activity, supresses *PhANG* (e.g. *LHCB2.1*) expression, and impairs carotenoid biosynthesis (MAYFIELD and TAYLOR, 1984; Brausemann et al., 2017; Dhami et al., 2022). NFZ blocked biosynthesis of cyclic carotenoids in etiolated seedlings, leaving only trace levels of lutein, zeaxanthin, and β-carotene discernible from residual seed-derived pigments (Figure S3A). De-etiolated NFZ-treated tissues appeared bleached without any appearance of viable photosynthetic chloroplasts (Figure S3B). NFZ treatment of WT etiolated seedlings significantly reduced the expression of *LHCB2.1* (94%), *RBCS1* (65%), and *εLCY* (65%) relative to untreated controls (Figure 1E). NFZ treatment did not alter the reduction in *εLCY* expression exhibited by *ccr2* (Figure 1F). NFZ treatment did block ζ-carotene, neurosporene and tetra-*cis*-lycopene biosynthesis in *ccr2* and caused a 4-fold hyperaccumulation of phytoene (and phytofluene) like that observed in WT (Figure S3B). Therefore, NFZ treatment of etiolated seedlings suppresses cyclic carotenoid biosynthesis and a feedback signal that negatively regulates *εLCY* transcript levels.

### Light-enhanced *εLCY* mRNA levels are suppressed in de-etiolated *ccr2* seedlings

We investigated if the expression of *εLCY* was altered during seedling photomorphogenesis and leaf development by comparing etiolated vs. de-etiolated cotyledon tissues and younger vs. fully expanded mature leaf tissues, respectively, from WT and *ccr2*. Relative to etiolated seedlings, the expression of *εLCY* significantly increased after 24 hrs in de-etiolated WT seedlings and there was 4.5-fold higher *εLCY* expression 72 hrs after photomorphogenesis (Figure 1G). Intriguingly, the *εLCY* expressions were consistently reduced by 2 to 2.5-fold across all time points in both etiolated and de-etiolated *ccr2* seedlings compared to WT (Figure 1G). The cyclic carotenoid content gradually increased in *ccr2* after 72 hours of illumination due to photoisomerization reaching less than 50% of the level displayed by WT (Figures S4A). Therefore, light induces *εLCY* transcript and cyclic carotenoid levels during photomorphogenesis.

Bioinformatics analysis using the Arabidopsis TAIR eFP browser revealed that the absolute expression of *εLCY* was strongest in younger rosette leaves (Figure S4B). We confirmed that *εLCY* transcript levels were 4-fold higher in younger relative to older WT leaves (Figure 1H). Young emerging leaves (leaf #10-13) contained higher carotenoid (and chlorophyll) levels compared to older fully expanded leaves (leaf #3-4) and do not normally accumulate linear *cis*-carotenes (Figure S4C-D). Therefore, *εLCY* expression and cyclic carotenoid levels decline during leaf development. Taken together, *εLCY* expression is negatively regulated by *ccr2* and in NFZ treated etiolated seedlings, declines during leaf development, and becomes positively regulated during light-mediated photomorphogenesis (Figure 1I).

### The *εLCY* promoter modulates metabolic feedback, developmental and light-mediated regulation of reporter gene expression

To decipher the feedback mechanism(s) regulating *εLCY* expression, an upstream fragment (- 450bp) of the *εLCY* promoter (plus the 5′UTR) was fused to the *FiLUC* reporter gene (*εLCY*Prom-*εLCY*5′UTR::*FiLUC*) and transformed into WT. Mature rosette leaf tissues from 23 independent F_2_ transgenic WT lines emitted low luminescence ranging from 295 to 7661 relative light units (RLU; Av. 3511) (Figure S5A), which was 17-fold lower than that enabled by the strong and constitutive CaMV35S promoter (Av. 59767 RLU; 37 *CaMV35S*::*FiLUC* independent F_2_ lines ranged from 480 to 220692 RLU) (Figure S5B) (Cazzonelli et al., 2010). *In planta* imaging of luminescence from two representative homozygous lines (WT-*εLP::FiLUC*#2e and WT-*εLP::FiLUC*#4c) revealed the *εLCY* promoter was active in the rosette leaves, cauline leaves, and petiole, but not in the roots (Figure 2A). The WT-*εLP::FiLUC* lines displayed lower luminescence in the fully expanded older leaves and stronger luminescence in the outer expanding regions from newly emerged leaves (Figure 2A). In contrast, the *CaMV35s* promoter enabled strong constitutive luminescence of all tissue types, including roots (Figure 2B). The transcript levels of *εLCY* and *FiLUC* in two *εLP::FiLUC* transgenic lines were 4- and 8-fold more abundant in younger (leaves 1-5) relative to older (leaves 6-10) leaves from mature rosettes, respectively (Figure 2C). FiLUC activity was, similarly, 5- to 7-fold higher in younger compared to older leaves from both *εLP::FiLUC* transgenic lines (Figure 2D). Therefore, the - 450bp *εLCY* promoter fragment is responsive to changes in development, driving higher *FiLUC* levels in younger compared to older leaves that contain less cyclic carotenoids.

**Figure 2.**
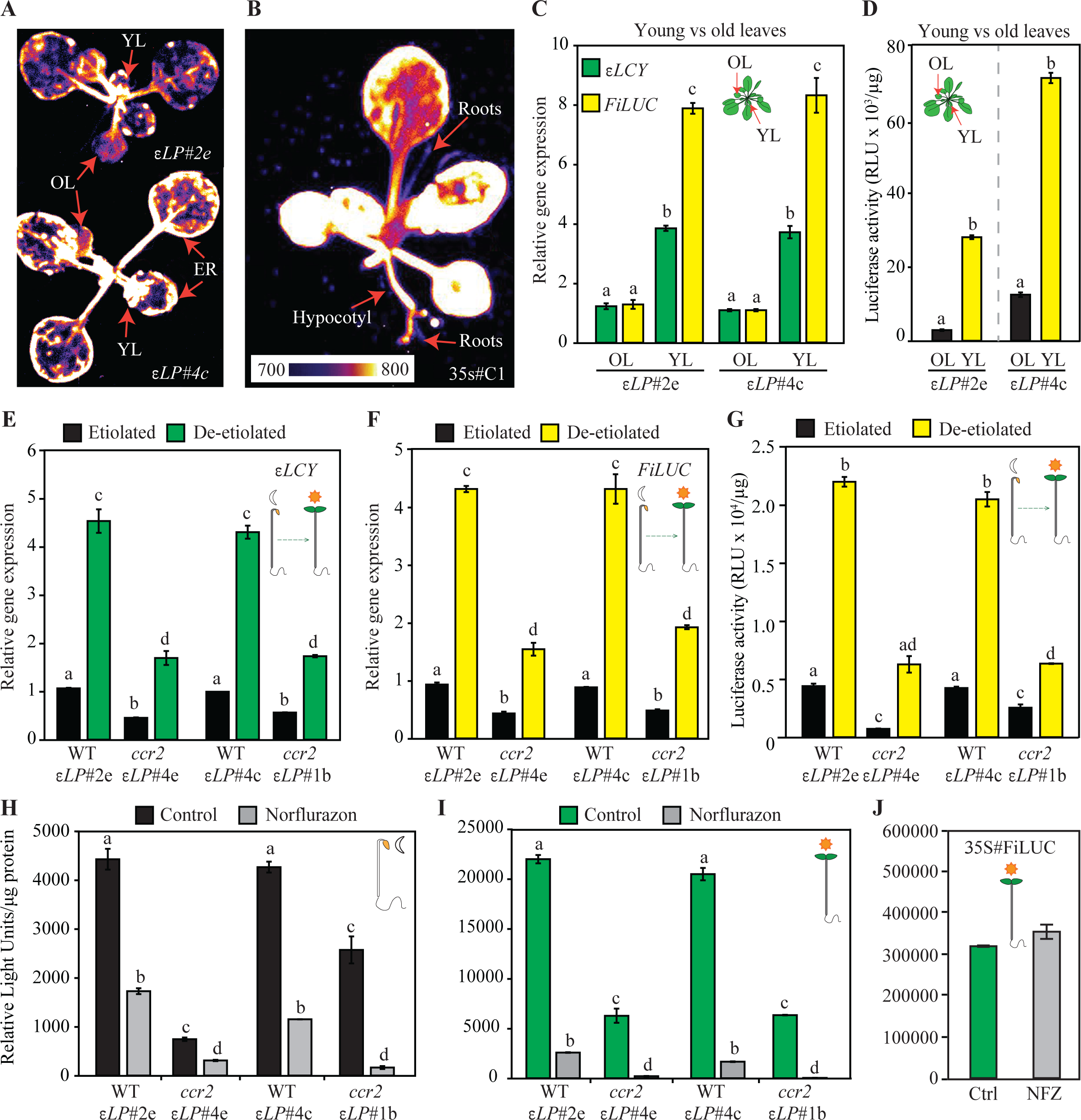
Quantification of *εLCY* promoter enabled *LUCIFEREASE* expression and reporter activity in response to chemical, genetic, developmental, and environmental changes. **A-B**) *In planta* luminescence emitted by independent WT transgenic plants harbouring the *εLCY*Prom-*εLCY*5′UTR::FiLUC (*εLP::FiLUC*#2e; *εLP*#2e, *εLP::FiLUC*#4c; *εLP*#4c) or CaMV35s::FiLUC (35S#C1) transgenes. The colour bar displays the intensity scale where black denotes no light emission and white indicates maximum luminescence. **C**) *εLCY* and *FiLUC* gene expression levels in young leaves (YL) and old leaves (OL) from WT transgenic lines (*εLP*#2e and *εLP*#4c). **D)** FiLUC activity (RLU/μg protein) in young and old leaves from WT transgenic lines (*εLP*#2e and *εLP*#4c). **E-G**) *εLCY* (E) and *FiLUC* (F) mRNA levels, and FiLUC activity (G) in transgenic etiolated and de-etiolated transgenic WT (*εLP*#2e, *εLP*#4c) and *ccr2* (*εLP*#4e, *εLP*#1b) seedlings. **H-I**) FiLUC activity in 5 μM NFZ-treated etiolated (H) and de-etiolated (I) transgenic WT (*εLP*#2e, *εLP*#4c) and *ccr2* (*εLP*#4e, *εLP*#1b) seedlings. (**J**) FiLUC activity in 5 μM NFZ-treated de-etiolated transgenic WT seedlings harboring 35S#C1. Lettering denotes significance by a one- or two-way ANOVA statistical analysis with post-hoc Tukey test. Data is representative of two to three independent experiments, and standard error bars are shown (n=3-9). Abbreviations: RLU; Relative light units, FiLUC; Firefly intron-containing LUCIFEREASE gene.

We next examined if the *εLCY* promoter fragment was responsive to metabolic feedback regulation triggered by NFZ-treatment of seedlings that lack biosynthesis of cyclic β-branch carotenoids in etiolated tissues (Figure S3A) and *ccr2* that shows delayed chlorophyll biosynthesis in de-etiolated seedlings (Cazzonelli et al., 2020). The WT-*εLP::FiLUC*#2e and WT-*εLP::FiLUC*#4c transgene lines were introgressed with *ccr2.1* (*ccr2.1*-*εLP::FiLUC*#4e) and *ccr2.5* (*ccr2.5-εLP::FiLUC*#1B) respectively, to create homozygous isogenic lines. Compared to WT, *εLCY,* and *FiLUC* gene expression were significantly reduced (>50%) in etiolated *ccr2* seedlings (Figure 2E-F). Three days after constant illumination, de-etiolated WT-*εLP::FiLUC* seedlings displayed a >4-fold increase in FiLUC as well as *εLCY* expression (Figures 2E-F). Etiolated and de-etiolated seedlings harbouring the *CaMV35S*::*FiLUC* transgene displayed similar *FiLUC* mRNA levels (Figure S5C). The *εLCY* and *FiLUC* expressions were approximately 50% lower in *ccr2*-*εLP::FiLUC* etiolated and de-etiolated seedlings relative to their WT isogenic parental line (Figure 2E-F). The fold changes in FiLUC activity enabled by the *εLCY* promoter in WT relative to *ccr2* and etiolated versus de-etiolated seedlings mirrored the *FiLUC* mRNA expression patterns (Figure 2F-G). The *εLP::FiLUC* promoter-reporter fusion was significantly repressed in both WT and *ccr2* etiolated and de-etiolated seedlings treated with NFZ, unlike CaMV35-FiLUC that showed no response to NFZ (Figure 2H-J). In summary, the *εLCY* promoter plus 5′UTR expression platform was responsive to *ccr2*-mediated feedback, NFZ inhibition of cyclic carotenoids, developmental changes in leaf aging, and light-mediated regulation during photomorphogenesis.

### The *εLCY* promoter harbours alternative transcription start sites and a conserved IRES element

*In-silico* analysis of *εLCY* promoter fragment (-450 bp) identified two TATA-like boxes (-305 bp and -357 bp) and CAAT boxes (-234 and -434 bp) located outside the consensus distances (-20 to -60 bp) defined for plant RNA Polymerase II promoters (Kumari and Ware, 2013; Shahmuradov et al., 2017) (Figure S6A). Conserved *cis*-elements such as the G-box (-213 bp), ABRE (-444 bp), auxin (TGA; -316 bp), and a light-responsive TCT-motif (-279 bp) were also found in the distal region of the promoter. 5′RACE analysis of the *εLCY* mRNA from etiolated (dark-grown) and de-etiolated (dark to light shift) WT seedlings was used to identify three 5’RACE PCR fragments (Figure 3A). Sequencing these products revealed three distinct alternative transcription start site ranges in dark-grown seedlings with a few outliers; TSS_1_ (-200 to -205 bp; < 218 bp), TSS_2_ (-130 to -143 bp; < 160 bp) and TSS_3_ (-70 to -75 bp; > 51 bp) (Figure 3B). TSS_1_ generated the longest 5′UTR fragment length of around -200 bp, specific to dark-grown seedlings. TSS_2_ and TSS_3_ generated two 5′UTR fragments (-133 bp and -70 bp) in dark and light-grown seedlings, with TSS_3_ being less abundant. The -133 bp 5′UTR fragment beginning at TSS_2_ was the more abundant 5’RACE product. TSS_2_ aligned with the transcription start of three EST cDNA transcripts deposited into GenBank (BP866039, BX831396 and EH842927). Sequence alignment of -234 bp of the *εLCY* 5’UTR from closely related *Brassicaceae* species (*A. thaliana, Arabidopsis lyrate*; *A. lyrata*, *Eutrema salsugineum*; *E. salsugineum*, *Capsella rubella*; *C. rubella*, and *Camelina sativa*; *C. sativa*) revealed similar sequence regions (referred to as a conserved domain; CD) in between and neighbouring the three *εLCY* transcription start sites (Figure 3C). That is, CD-1 (-234bp to -213 bp) was positioned upstream of TSS_1_, while CD-2 (-190 to -155 bp) was located downstream in between TSS_1_ and TSS_2_. CD-3 (-124 to -87 bp) was located downstream of TSS_2_ prior to TSS_3_ (Figure 3B). Therefore, the *εLCY* promoter does not appear to have a canonical TATA box, yet generates a dominant -133 bp 5′UTR fragment in etiolated and de-etiolated seedlings.

**Figure 3.**
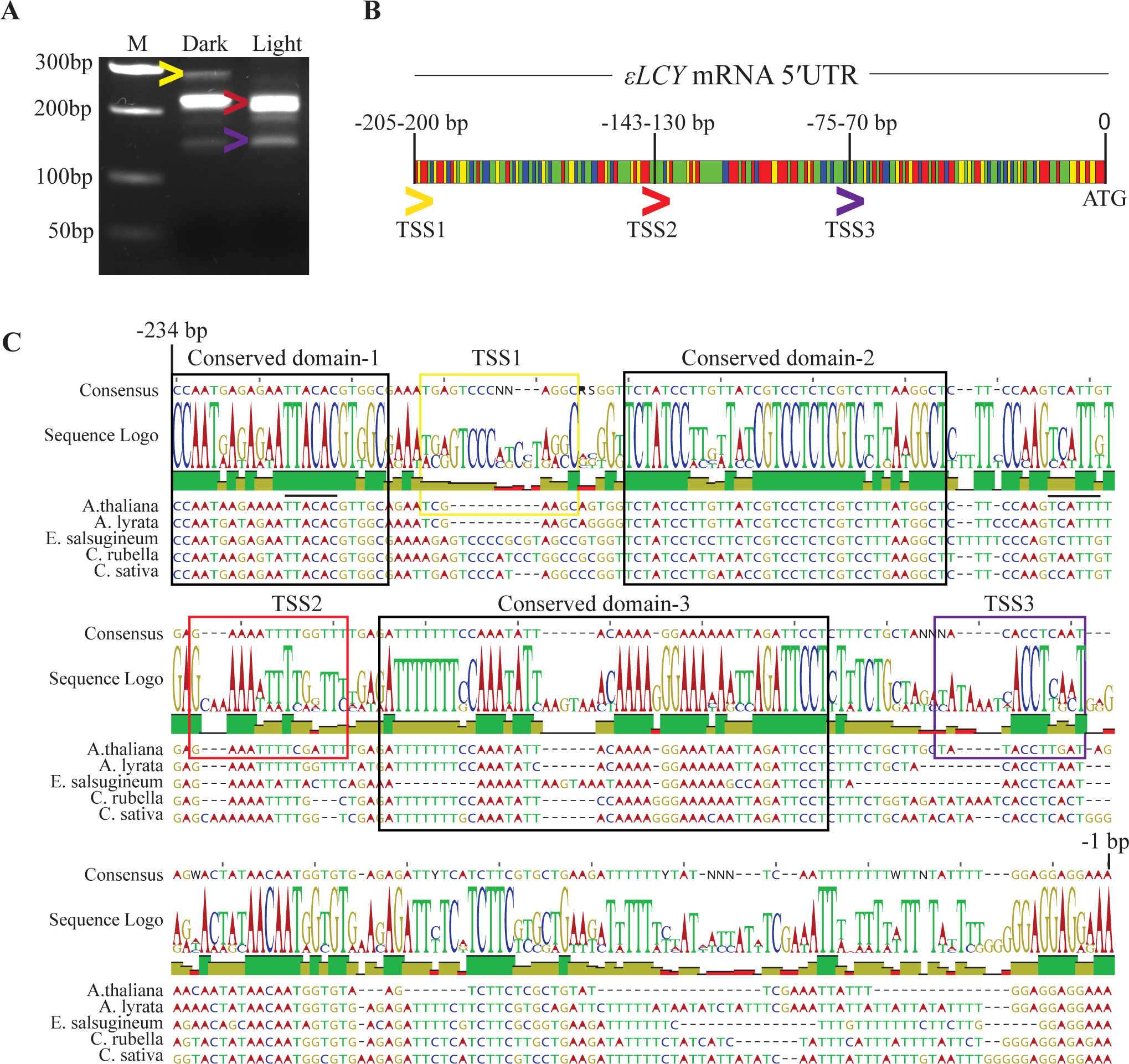
Analysis of *εLCY* mRNA transcription start sites and 5′UTR sequence homology among *Brassicaceae* species. **A**) Gel electrophoresis of 5′RACE fragments of the *εLCY* 5′UTR in etiolated (dark) and de-etiolated (light) grown wild-type seedlings. Coloured arrowheads denote the 5’RACE PCR fragments on the gel electrophoresis image that are representative of at least three independent 5’RACE experiments. **B**) Sequence representation of the three alternative transcription start sites (TSS) positioned around -200, -130, and –70bp upstream from the translational start codon sequence (ATG). The coloured arrowheads depicted below the sequence reflect three alternative 5’UTR lengths determined by PCR, and the identifier bar depicts the nucleotide composition (Adenine; red, Thymine; green, Guanine; yellow, Cytosine; blue). **C**) *εLCY* 5′UTR sequence showing three alternative TSS positioned proximal to evolutionary conserved sequence domains (black box) identified among *Brassicaceae* species. Sequences of *Arabidopsis thaliana* (*A. thaliana*; NC_003076), *Arabidopsis lyrata* (*A. lyrata*; NW_003302548), *Eutrema salsugineum* (*E. salsugineum*; NW_006256829), *Capsella rubella* (*C. rubella*; NW_006238916) and *Camelina sativa* (*C. sativa*; NC_025702) were aligned using Geneious Prime Software (v10.2.6). NCBI identifiers of sequences analysed are shown above in brackets. The two black lines above the *A. thaliana* sequence (3-10 nucleotides upstream of TSS_1_ and TSS_2_ regions) denote homology to the consensus Initiator (Inr) element (Figure S5).

Transcription of TATA-less promoters can be determined by other core-promoter elements, such as the Initiator element (Inr), downstream promoter element (DPE), and the pyrimidine patch (YP), all of which align with the three *εLCY* transcription start sites (Figure S6A). Inr sequence elements (Py-Py-A-N-T/A-Py-Py; where Py is C/T, and N is A,G,C,T) promote transcription initiation in plants by acting as recognition sites to bind TFIID (Yamamoto et al., 2007; Srivastava et al., 2014). Two Inr motifs identified were positioned 3-10 nucleotides upstream from TSS_1_ (-222 bp) and TSS_2_ (-153 bp) that aligns closely with the consensus range (- 14 to +14bp) defined for plant TATA-less promoters (Figure 3C, S6A) (Yamamoto et al., 2009; Shahmuradov et al., 2017). Two DPE elements (-256 bp, -343 bp) reported to function with Inr to bind TFIID were identified upstream from TSS_1_ yet are located outside the consensus distribution (+25 to +35 bp) relative to the TSS of plant dicot promoters (Kumari and Ware, 2013). The YP is a core promoter element usually located upstream from the TSS (-50 bp) and downstream from the TATA or INR motifs. Two YP motifs were identified -44 bp (-179 bp) and -11 bp (-86 bp) upstream from TSS_2_ and TSS_3,_ respectively, which generate the two main *εLCY* 5’UTR fragments on either side of CD-3 (Figure S6A). A third YP was identified (-37 bp) immediately downstream of a putative upstream open reading frame (uORF) (Met-Val-STOP; - 47 bp to -39 bp) within the *εLCY* 5’UTR (Figure S6A). An intron-mediated enhancer (IME)-like motif (TTNGATTTG; 88% homology) harbouring the pentamer core sequence “CGATT” located at -137 bp overlapped with TSS_2_. IME can enhance transcription and is often enriched in 5′UTRs near the TSS, where it is proposed to signal an increase in RNA polymerase processivity (Parra et al., 2011; Laxa et al., 2016; Gallegos and Rose, 2017). Finally, a conserved internal ribosome entry site (IRES) structural motif common to RNA viruses was identified using UTRscan to span the 5′UTR (-1 to -82 bp) starting downstream of CD-3 and overlapped two YP motifs that separate TSS_3_ (-86 and -37 bp) (Figure S6A). IRES elements can adopt structural configurations that function to recruit ribosomes, initiation factors, and/or RNA-binding proteins (Etzel and Mörl, 2017; Martinez-Salas et al., 2018). The proximal location of IME, Inr, DPE, and YP motifs relative to the transcription start sites could link transcriptional enhancement with IRES-mediated translational regulation via an RNA structural rearrangement of the *εLCY* 5’UTR.

### The *εLCY* 5’UTR does not impact FiLUC activity enabled by the CaMV35S promoter

Variations in the *εLCY* 5’UTR were shown to modify β-carotene levels in maize and wheat (Richaud et al., 2018). This prompted us to further investigate a potential function for the *εLCY* 5′UTR in modulating downstream expression. A gain-of-function approach was used to interrogate the *εLCY* 5′UTR fragment (starting at -133 bp, TSS_2_ identified by 5′RACE) by testing mutation and deletion (MutDel) versions placed in between the CaMV35s promoter and the *FiLUC* reporter gene. The CaMV35S promoter served as the control (Ctrl-35S) and MutDels of the *εLCY* 5′UTR tested potential functions of 1) the upstream open reading frame (-uORF; putative internal 3 amino acid peptide), 2) the out-of-frame uORF with the main coding sequence (-ATG), and 3) the IRES (-5′IRES and -3′IRES) (Figure 4A). Luciferase activity was quantified in foliar tissues from independent Arabidopsis transgenic lines (Figure 4B) and agro-infiltrated tobacco plants (Figure 4C) ectopically and transiently expressing the promoter-reporter fusions, respectively. The -133 bp *εLCY* 5’UTR fragment had no significant effect on FiLUC activity relative to the Ctrl-35S lacking any 5’UTR. Removal of the small three amino acid uORF and mutation of the initiation start codon (-ATG) that would prevent a putative out-of-frame uORF from the *εLCY* 5’UTR had no significant impact on FiLUC activity. Removal of CD-3 and evaluation of the TSS_3_ fragment did not affect luciferase activity relative to the -133bp *εLCY* 5’UTR. Deletion of the 5’/3’ ends of the IRES element similarly had no significant effect on FiLUC activity in transgenic or tobacco leaves with transient expression (Figure 4B-C). In summary, mutations and deletions that alter 5’UTR length, remove an uORF, and structural IRES motif did not affect CaMV35s enabled luciferase activity.

**Figure 4.**
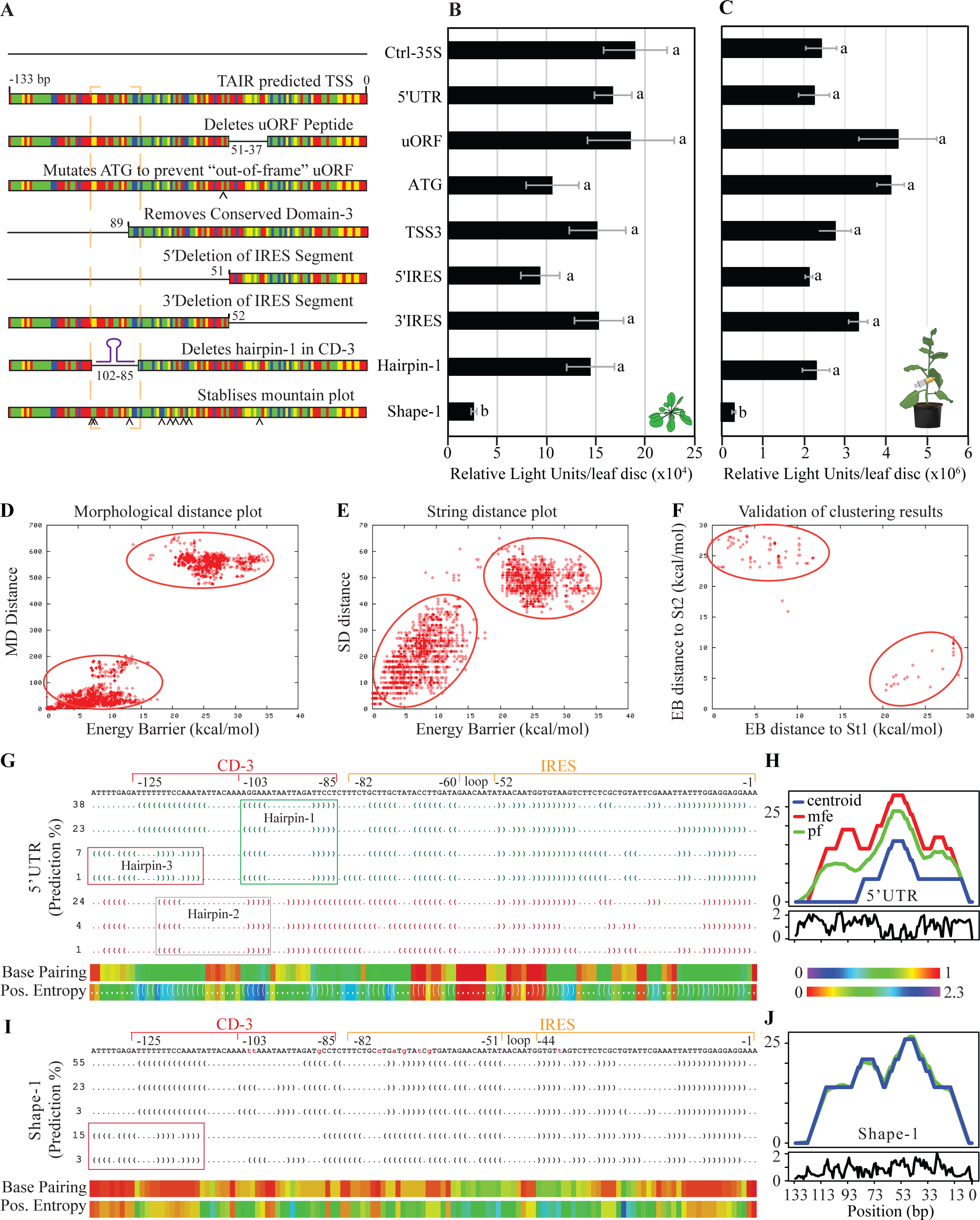
RNA structural analysis of the *εLCY* 5′UTR and regulation of promoter-reporter gene activity in transgenic Arabidopsis and transiently-infected tobacco leaves. **A**) Diagrammatic representation of the various synthetic mutations and deletions (MutDels) of the *εLCY* 5′UTR fused in between the CaMV35S promoter and *FiLUC* reporter gene. Constructs tested include; Ctrl-35S (Control CaMV35S promoter with no 5′UTR), 5′UTR (5’RACE and TAIR10 predicted TSS; -133 to -1bp), uORF (removes highly conserved sequence and small uORF; -51 to -37 bp, ATG (mutates ATG start codon to prevent formation of a putative out-of-frame uORF, TSS_3_ (Includes entire TSS_3_ fragment and removes CD-3; -90 to -1 bp), 5′IRES (Removes 5′ segment of IRES and CD-3; -133 to -51 bp), 3′IRES (Removes 3′ segment of IRES; -52 to -1 bp), Hairpin (deletes stem-loop structure in CD-3; -102 to -85 bp), and Shape-1 (9 serial substitution mutations that disrupt structural switching). Colour bars represent nucleotides (A; red, T; green, C; blue, G; yellow). **B**) Average Relative lights units (RLU) in leaf tissues from transgenic Arabidopsis plants harbouring Ctrl-35s and MutDel transgenes. Equal-sized leaf discs (two discs per leaf) were dissected from two leaves from multiple plants (4-10) and transgenic lines (4 x Ctrl-35S, 6 x 5’UTR, 12 x uORF, 10 x Hairpin, 9 x ATG, 5 x TSS_3_ 8 x 5’IRES, 9 x 3’IRES, 7 x Shape-1). **C**) Average Relative Lights Units (RLU) emitted by agrobacteria-infiltrated tobacco leaf tissues transiently expressing promoter-reporter fusions. A leaf disc bioassay (n=2 leaf discs per plant) from multiple plants (n=8-12) was used to quantify luciferase activity. Data represents the average of three independent experiments (n=3). Luminescence (relative light units) was quantified using an *in vivo* luciferase activity assay (Cazzonelli and Velten, 2006). **D-F**) Structural conformational switch prediction of -133bp of the *εLCY* 5′UTR using paRNAss software. Morphological distance plot (A), string distance plot (B), and validation of clustering results (C). The energy required to convert different RNA structures into each other is given with the energy barrier (EB; in kcal mol^-1^). The alignment distance is a measure of the structural similarity of the predicted structures to convert between different RNA structures. The cluster of each kinetically related structure with the same consensus folding conformation forms a separate energy cloud. There must be at least two energy clouds with distinct separation on the x- and y-axis of the distance plot to indicate any switching potential. **G, I)** RNAShapes analysis of the 5′UTR (G) and shape-1 (I) sequences showing Vienna shape representatives (dot-bracket notation; ’(’ and ’.’ represent paired and unpaired bases respectively), probability prediction (%), base-pairing probability colour chart (1 implies highly stable), and positional entropy colour chart (0 indicates base-pair is always unpaired or paired with its partner) of the most probable structures generated by RNAfold software default settings. Green or red dot-bracket notations denote similar representations. Red and orange lines above the *εLCY* 5’UTR sequence denote Conserved Domain-3 (CD-3) and IRES motif. Boxed regions denote hairpin structures. **H, J**) RNAfold mountain plot representations showing the thermodynamic ensemble of RNA structures of the *εLCY* 5’UTR (**H**) and shape-1 (**J**) sequences. "mfe" represents minimum free energy structure; "pf" indicates partition function; "centroid" represents the best average structure in a plot of height m(k) versus position (bp), where the height is given by the number of base pairs enclosing the base at a position. Loops correspond to plateaus (hairpin loops are peaks), and helices to slopes. Entropy of each bp along the RNA sequence (-1 to -133 bp) is shown below the mountain plot and lower entropy values indicate higher stability of RNA secondary structures.

### RNA structural switching within the *εLCY* 5’UTR suppresses FiLUC activity

To investigate the possibility of RNA secondary structural rearrangements modulated by the IRES, we analysed the -133 bp (TSS_2_) *εLCY* 5′UTR fragment using RNA structural definition softwares, namely paRNAss (Giegerich et al., 1999; Voss et al., 2004; Álvarez et al., 2016), RNAShapes (Steffen et al., 2006), and RNAfold (Gruber et al., 2008; Lorenz et al., 2011). paRNAss interrogated the folding space for alternate RNA secondary structures within an energy range above the MFE structure. Two distinct and separate energy clouds were displayed on plots of morphological distance, string distance, and validation of clustering indicating energetically separable RNA structural populations - one near the x-axis and one near the y-axis, indicating some potential to switch between structural configurations (Figure 4D-F). RNAShapes revealed seven RNA structural predictions (>1% probability) having a minimal free energy (mfe) ranging from -19.1 to -22.4 kcal/mol. There were two major RNA configurations, one dominating 38% (green font) and the alternative 24% (red font) of probable structural classes that had a central loop (-52 to -60 bp) within the IRES motif (-1 to -82 bp) of all structural predictions and were distinguished by hairpin configurations within CD-3 (Figure 4G, S7A). The RNAfold mountain plot representation in a plot of height versus position confirms a prominent centroid structural peak (-58 bp) within the IRES that was surrounded by a higher probability of base-pairing and lower minimal positional entropy indicative of fewer secondary structure outcomes within this probability space (Figure 4G-H). In contrast, the CD-3 region lacked a centroid structure, the mfe and partition function (pf) structural predictions were displayed as separate peaks, there was a reduced probability of base-pairing and higher positional entropy surrounding the predicted hairpin structures (Figure 4G-H). Hairpin-1 (-85 to -103 bp) and hairpin-2 (-98 to -120 bp) were distinguishable in the four dominant and three alternative structural classes, respectively representing 69% and 29% of all probabilities. Whereas hairpin-3 (-97 to -133 bp) was identified in only two of the four dominant structural probability configurations representing 8% of structural predictions (Figure 4G, S7). Taken together, there appears to be RNA structural similarity within IRES, and three alternative hairpin configurations within CD-3.

To determine if alternative RNA structural configurations can regulate reporter activity, nine serial mutations were introduced into the 5′UTR sequence to generate a stable mountain plot representation with negligible separation between the structure prediction curves (centroid, partition function; pf, and mfe) (Figure 4J). All structural predictions retained the IRES central loop (-44 to -51 bp) that was slightly offset compared to the 5′UTR (Figure 4I). A single RNA shape (shape-1) dominated 81% of structural predictions having similar dot-bracket notations within the IRES region and a higher base pairing probability of binding with lower positional entropy within CD-3. This led to the removal of hairpin-1 and stabilization of hairpin-3 in 18% of structural probabilities (Figure 4I, S7). Intriguingly, the shape-1 5′UTR fragment when inserted between the CaMV35s promoter and luciferase reporter led to the suppression of FiLUC activity (> 3-fold) in transgenic Arabidopsis leaves as well as in tobacco leaves infiltrated with agrobacteria harbouring the promoter-reporter fusion (Figure 4A-C). Removal of the hairpin-1 sequence from CD-3 (-85 to -103 bp) did not affect CaMV35s enabled luciferase activity in leaves (Figure 4A-C). Therefore, 5′UTR serial mutations predicted to stabilize the RNA structure have imparted a negative regulating expression platform.

### Stabilisation of the 5’UTR hairpin-3 structure promotes posttranscriptional regulation

The mechanism regulating the negative expression platform mediated by shape-1 was further examined in mature leaves, etiolated and de-etiolated seedlings. The *FiLUC* mRNA to activity ratio was quantified in mature leaf tissues from three independent transgenic lines harboring the 5′UTR and shape-1 fragments relative to a 35s promoter control (no 5′UTR; exhibits similar reporter activity to the 5′UTR) (Figure 5). The average ratio of three lines haboring shape-1 (9.4-fold) were significantly higher than the average ratio of three 5′UTR lines (2.7-fold) due to lower activity levels relative to similar transcript levels (Figure 5A). In etiolated and de-etiolated seedling tissues, the reporter activity levels were also suppressed (5.2- to 2.4-fold) and yet transcript levels remained high (2.3- to 6-fold) in shape-1 (#1-4) relative to 5′UTR (#7A2) (Figure 5B-C). Agrobacteria-mediated transient expression in *N. tabacum* leaves was used to circumvent any transgene chromosomal position effects influencing promoter-reporter regulations in independent transgenic lines. CaMV35s, 5’UTR, and shape-1 displayed similar *FiLUC* transcript levels, while FiLUC activity was severely suppressed by shape-1 (9- to 15-fold; Figure 5D-E). Therefore, shape-1 imparts a posttranscriptional expression platform that suppresses CaMV35 promoter enabled luciferase activity *in planta*.

**Figure 5.**
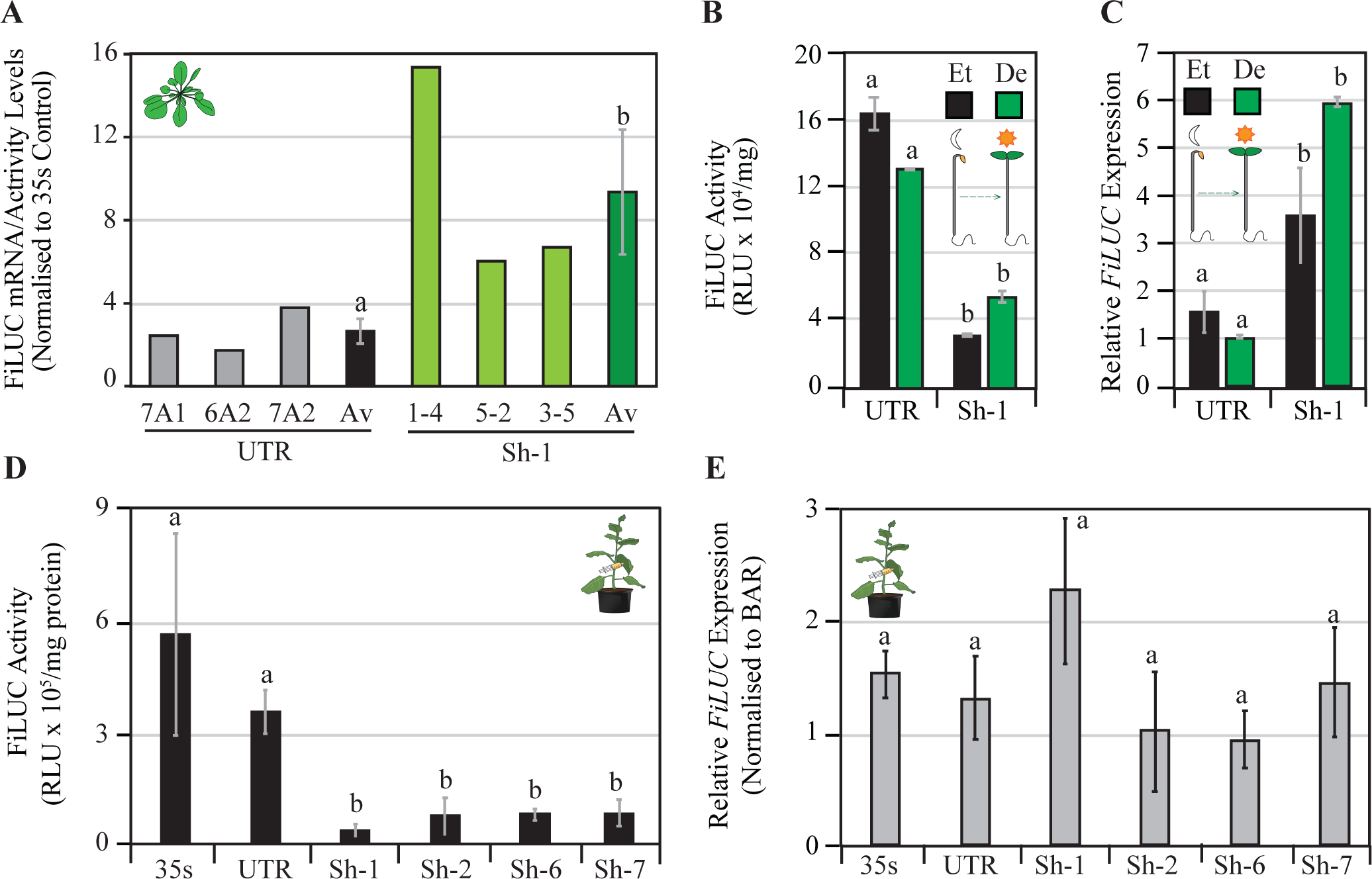
Regulation of LUCIFERASE activity and mRNA levels by the *εLCY* 5′UTR and shape fragments in etiolated, de-etiolated and leaf tissues. **A**) Ratio of FiLUC mRNA to activity (RLU/mg protein) in mature leaves from transgenic Arabidopsis lines harboring UTR and shape-1 (Sh-1) fragments normalised to the CaMV35s promoter control. Data is representative of a single experiment showing the average mean and standard error of three independent lines (n=3). **B-C**) FiLUC activity (RLU/mg protein) (B) and relative *FiLUC* mRNA expression levels (C) in etiolated (Et) and de-etiolated (De) transgenic Arabidopsis seedlings harbouring the UTR and shape-1 fragments. Data is representative of two experiments showing the mean and standard error (n=3). **D-E)** Quantification of FiLUC activity (RLU/mg protein) (D) and relative *FiLUC* mRNA levels (E) in agro-infiltrated tobacco leaf tissues. The *BAR* (Basta-resistance) gene was used to normalise between agro-infiltrations, and *ACTIN* served as an internal housekeeping gene for qRT-PCR normalisation. Promoter constructs tested include CaMV35s (*35S*), CaMV35s-*εLCY*5′UTR (UTR), CaMV35s-Shape-1 (Sh-1), CaMV35s-Shape-2 (Sh-2), CaMV35s-Shape-6 (Sh-6), CaMV35s-Shape-7 (Sh-7). Data represents the average of two independent experiments and six biological replicates with standard error bars shown (n=6). Lettering denotes significance (one-way ANOVA, p<0.05).

Mutations within shape-1 were sequentially restored back to their WT sequence to interrogate if RNA structural changes or the gain of a *cis*-acting motif functions to modulate the posttranscriptional expression platform (Figure S6B). Shape-2 (8 mutations), shape-6 (3 mutations) and shape-7 (2 mutations; -102 bp, GG > TT) were designed based upon having negligible separation between the structure prediction curves in their mountain plot representations (Figure S8A). The central loop region within the IRES element of the 5’UTR dot-bracket notations was restored in shape-6 and shape-7 and surrounded by a high probability of base pairing and lower entropy (Figure S8B-D). Shapes-2/6/7 were not predicted to form hairpin-1 (between -102 to -85 bp) but rather had a higher probability to form hairpin-3 (-97 to -133 bp) in CD-3 (Figure S7, S8B). Hairpin-3 was dominated in 85% of the structural definitions of shape-7 that harboured two mutations (GG>TT) disrupting hairpin-1 and hairpin-2 stem-loop formation (Figure S7E, S8B-D). In comparison to the CaMV35s and 5’UTR, shape-2 (8 mutations), shape-6 (3 mutations), and shape-7 (2 mutations) showed a significant reduction in luciferase activity (-4.3, -6.8, and 3.5-fold respectively) just like shape-1 when transiently expressed in *N. tabacum* leaf tissues (Figure 5D). Quantification of *FiLUC* mRNA abundance (normalised to internal BAR housekeeping transgene) in the agro-infiltrated tobacco leaf tissues from *N. tabacum* revealed that shapes-1/2/6/7 all had similar transcript levels to that of the 5’UTR and CaMV35s (Figure 5E). Therefore, disruption of hairpin-1 and hairpin-2 and stabilisation of a single RNA structural definition haboring hairpin-3 confers a negative regulating posttranscriptional expression platform.

### Shape-1 harbours an aptamer domain responsiveness to metabolic feedback

A command-line version of Infernal (1.1rc1, cmscan, default parameters) was used to scan the - 143 bp *εLCY* 5’UTR (AT5G57030; Chr5:23077255-23080053) against Rfam covariance model homology within the Rfam database collection for RNA families. The search stringency was reduced identifying some weak homology (E-value p<0.07) between an SMK_box riboswitch (RF01767; 81 bp) and the *εLCY* 5′UTR (-96bp to +13 bp of mRNA) fragment overlapping the IRES motif (Data File S1). We undertook a modelled approach to ascertain if the *εLCY* 5′UTR had other RNA structural characteristics similar to 27 different classes of metabolite-binding RNA riboswitches found in bacteria (Data File S2). Bacterial nucleic acid sequences retrieved from the Rfam database were analysed using RNAshapes software to ascertain structural features, including the number of shapes (between 1 to 5; p>0.1 or 10%), variants per shape, and minimum/maximum of shape forming probabilities (%) as well as minimal free energy (mfe; kcal) for 2691 RNA riboswitches (Table 1). Most riboswitches displayed one to two structural shape definitions (35% and 42%, respectively), and a considerable proportion (23%) displayed between three to five structures. The *εLCY* 5′UTR (-133 bp) displayed three structural definitions (38, 23 and 24% probability) ranging between the minimum (11%) and maximum (58%) probabilities displayed by 457 variants from 25 riboswitch classes (Figure 4G; Table 1). In comparison, shape-1 also generated three similar structural probabilities of 55, 23 and 15%, representing 93% of all predicted structural definitions. Therefore, the 5’UTR and shape-1 fragments can display structural features and a switching potential resembling metabolite-binding RNA riboswitches.

**Table 1.** RNAshapes modelling of structural features from 27 different classes of bacterial metabolite-binding RNA riboswitches.

| Number of shapes | Variants per shape | Shape Forming Probability (%) |  | Minimal Free Energy (kcal) |  |
| --- | --- | --- | --- | --- | --- |
|  |  | Minimum | Maximum | Minimum | Maximum |
| **1** | 948 | 77 | 100 | -81 | -6 |
| **2** | 1123 | 20 | 78 | -79 | -6 |
| **3** | 457 | 11 | 58 | -77 | -6 |
| **4** | 143 | 11 | 45 | -82 | -14 |
| **5** | 20 | 10 | 29 | -68 | -24 |

We tested if the 5′UTR and/or a Shape-1 fragments were responsive to metabolic feedback triggered in *ccr2* or NFZ-treated seedlings that have impaired biosynthesis of β-branch cyclic carotenoids (Figure S3A). De-etiolated WT transgenic seedlings harbouring the CaMV35s and 5′UTR promoter fragments showed little response to NFZ treatment displaying similar FiLUC activity and transcript levels (74.6% to 111.6%) relative to the untreated controls (Figure 6A-B). Surprisingly, the Shape-1 and the -450 bp *εLCY* promoter fragments displayed significantly lower FiLUC activity (28.6% and 25.6% respectively) and mRNA levels (12.7% and 30.6% respectively) compared the untreated controls, which was 4- to 7-fold lower than the 5′UTR. We next selected independent WT and *ccr2* transgenic lines harbouring the shape-1 fragment that displayed similar FiLUC activity levels in mature leaves from plants that do not exhibit carotenoid-mediated feedback when grown under a 16 h photoperiod (Figure 6C). The FiLUC activity levels of shape-1 in WT and *ccr2* genotypes were >5-fold lower than the 5’UTR and CaMV35s promoters as expected. In de-etiolated seedlings, FiLUC activity was markedly higher in WT compared to *ccr2* that displays metabolic feedback inhibition (Figure 6D). Regardless of the genotype (WT vs *ccr2*), shape-1 activities in de-etiolated seedlings were considerably lower than the CaMV35 and 5′UTR promoters. Therefore, the shape-1 fragment appears to harbour an aptamer domain that can sense metabolic feedback signalling and modulate changes in reporter gene expression.

**Figure 6.**
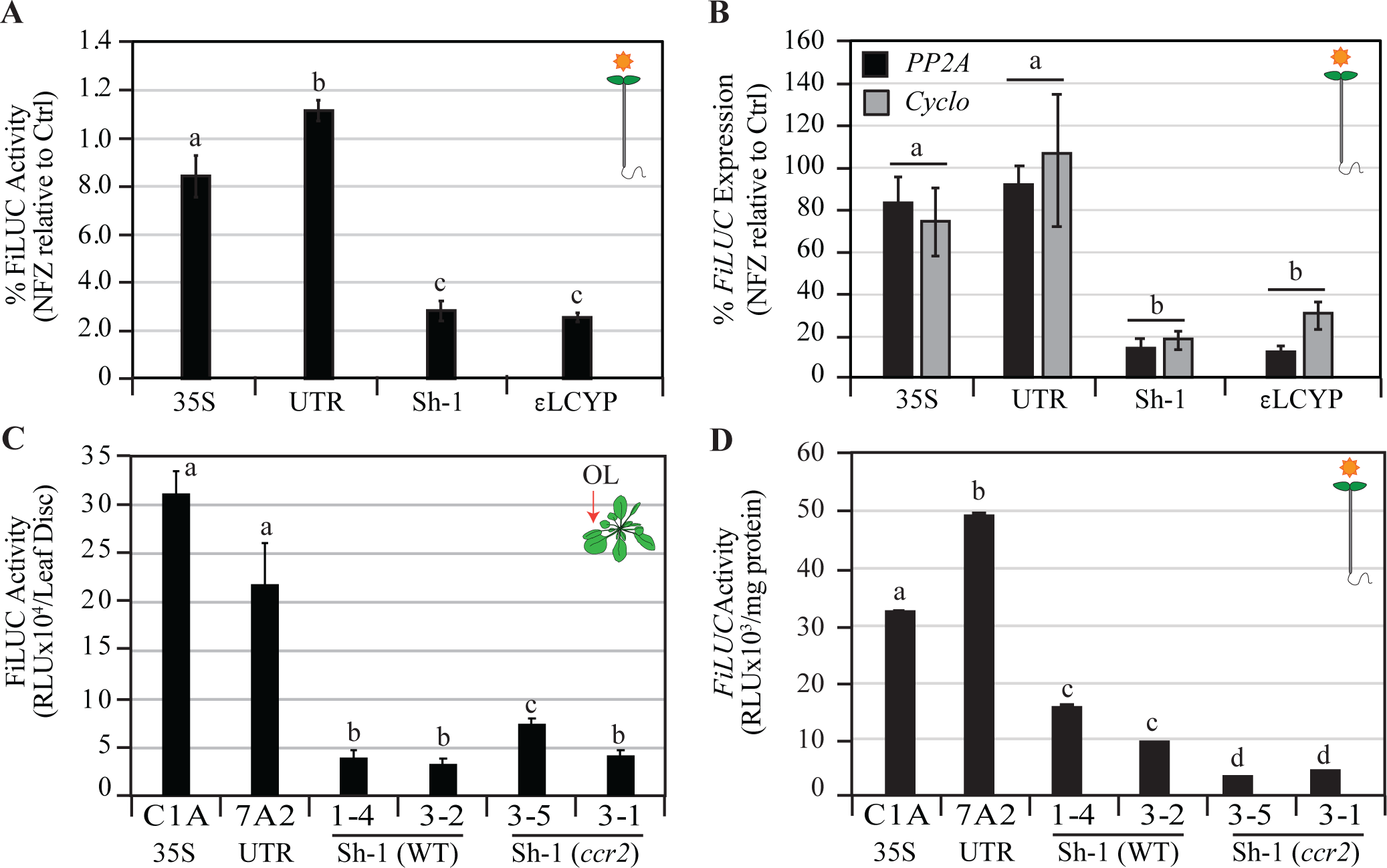
Norflurazon and *ccr2*-mediated feedback regulation of LUCIFERASE activity and mRNA levels by the 5′UTR shape-1 in de-etiolated tissues. **A-B**) Percentage (%) FiLUC activity (A) and *FiLUC* mRNA expression levels (B) in de-etiolated transgenic Arabidopsis seedlings grown on artificial media in the absence (Control; Ctrl) or presence of norflurazon (NFZ; 5μM). Percentages reflect the ratio of NFZ-treated relative to untreated control (Ctrl) seedlings. Transgenic lines harbour the 35S, UTR, Sh-1, *εLCY* (*εLCYP*) promoter-reporter gene fusions. **C-D**) Quantification of FiLUC activity (RLU/mg protein) in mature leaves (C) and de-etiolated seedlings (D) from WT and *ccr2* transgenic lines harbouring the Sh-1 fragment. Mean and standard error (n=3) are representative of two independent experiments. Lettering denote significance (one-way ANOVA, p<0.05).

## DISCUSSION

### Regulation of *εLCY* expression is not triggered by linear *cis*-carotene accumulation

The loss-of-function in *crtiso* (*ccr2*) triggers *cis*-carotene accumulation and suppresses *εLCY* expression in etiolated seedlings (Cuttriss et al., 2007). We recently reported a *cis*-ACS regulated plastid biogenesis, *HY5*, *PIF3*, and *PhANG* expression in etiolated and de-etiolated seedlings, as well as young emerging leaves of *ccr2* that accumulate linear *cis*-carotenes (Cazzonelli et al., 2020; Dhami et al., 2022). We questioned if metabolic feedback regulation of *εLCY* expression was also linked to linear *cis*-carotene accumulation. Etiolated cotyledon tissues from *ccr2*, *ziso*, *det1-1*, and WT treated with NFZ, all showed reduced *εLCY* expression that did correlate with the accumulation of linear *cis*-carotenes and a reduction in cyclic carotenoids. The loss-of-function in ZISO (*ziso-155*) and impaired activity of DET1 (*det1-154*) were previously shown to reduce linear *cis*-carotene levels in *ccr2* etiolated seedlings and restoring PLB formation and *PhANG* expression by limiting the production of a yet-to-be identified *cis*-ACS (Cazzonelli et al., 2020). When compared to WT, the *εLCY* expression remained suppressed in *ccr2 ziso* and *ccr2 det1-154* etiolated seedlings despite a reduction in di-*cis*-ζ-carotene, neurosporene and tetra-*cis*-lycopene which are proposed substrates of the *cis*-ACS. Therefore, the accumulation of linear *cis*-carotenes and a yet-to-be discovered *cis*-ACS does not appear to trigger metabolic feedback regulation of *εLCY* expression during skotomorphogenesis.

### Changes in cyclic carotenoids trigger metabolic feedback regulation of *εLCY* expression

Genetic manipulations, dark to light shifts, chemical treatments, and developmental change in Arabidopsis tissues showed a link between β-branch carotenoid and *εLCY* transcript levels (Table S1). NFZ treatment of seedlings blocks downstream carotenoid biosynthesis and elicits a retrograde signal that suppresses *PhANG* expression to thereby impair plastid biogenesis (Wu and Bock, 2021). *ccr2* also blocks carotenoid accumulation in etiolated tissues, yet the lack of any PLB formation in *ccr2* etioplasts and a unique transcriptomic profile indicated that *ccr2* could trigger both independent and overlapping signalling pathways (Cazzonelli et al., 2020; Dhami et al., 2022). While *εLCY* has not been considered a traditional *PhANG* like *LHCB1,* our evidence reveals that both these genes are repressed in NFZ-treated etiolated seedlings. Unlike NFZ-treated tissues, chloroplast biogenesis and carotenoid biosynthesis are delayed in *ccr2* de-etiolated seedlings as well as young emerging leaf tissues, both of which exhibit lower *εLCY* expression levels indicative of metabolic feedback regulation. The suppression of *εLCY* expression by RNAi or mutations redirects metabolic flux towards the β-branch of carotenoid biosynthesis pathway (Diretto et al., 2006; Harjes et al., 2008; Shi et al., 2015; Ke et al., 2019). Similarly, the exogenous expression of the *Pantoea ananatis* phytoene desaturase (CRTI) gene in wild-type, *tangerine, or old gold crimson* tomato increased β-carotenoid levels as well as *εLCY* expression during fruit ripening (Enfissi et al., 2017). Here we demonstrated exogenous expression of *PaCrtI* or the loss-of-function in CCD4 in *ccr2* etiolated tissues increased β-branch carotenoids (β-carotene and zeaxanthin) and induced higher *εLCY* transcript levels. It is yet to be determined if the presence or absence of a signalling molecule represses or promotes *εLCY* expression. A plastid-derived retrograde signal could feedback to modulate *εLCY* expression (Wu and Bock, 2021). The feedback regulation of *εLCY* expression is connected to changes in β-branch carotenoids, suggesting that a β-apocarotenoid signal may feedback to balance alpha and beta-branch carotenoid homeostasis during plastid biogenesis (Figure 7).

**Figure 7.**
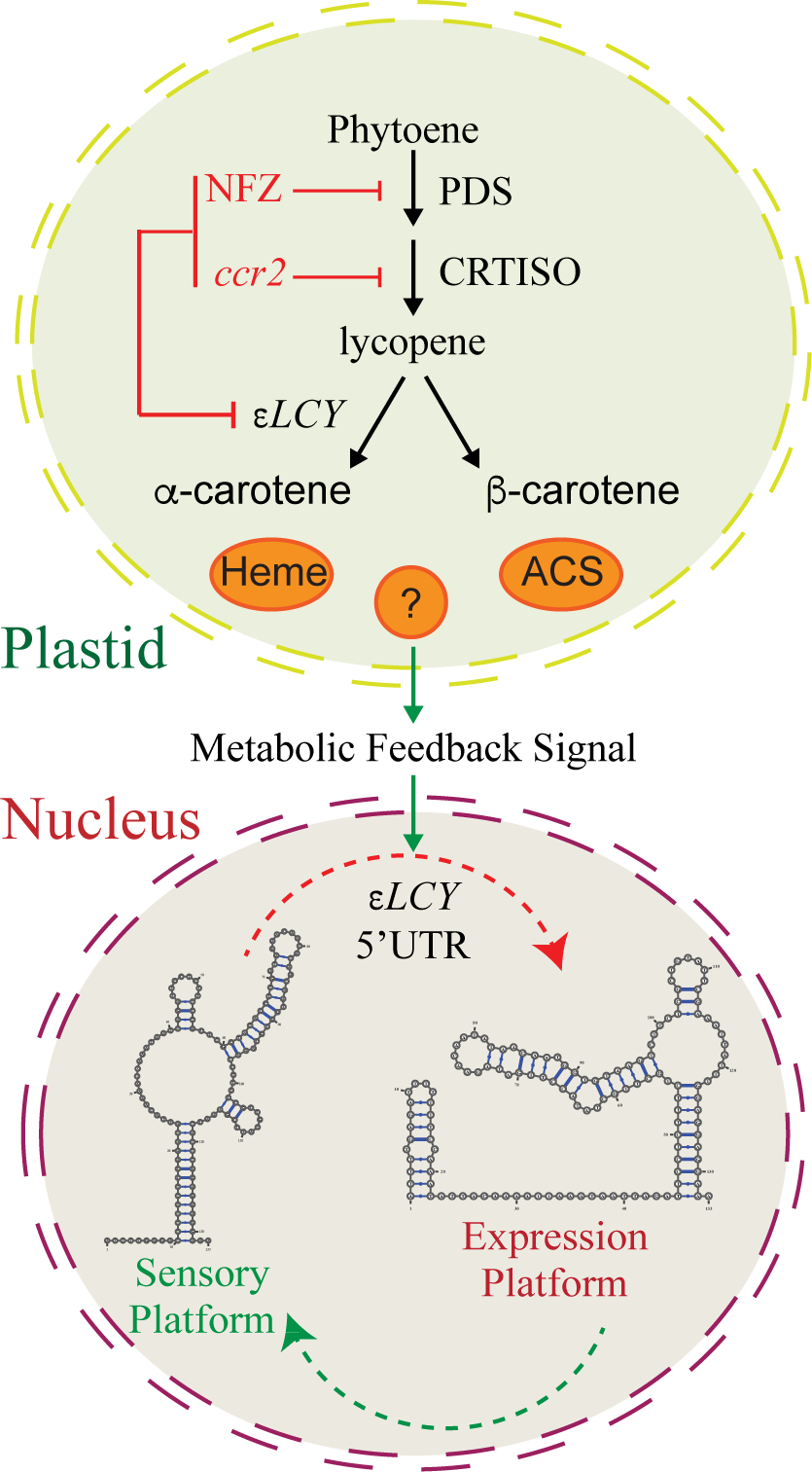
Proposed model showing carotenoid-mediated feedback signalling between the nucleus and plastid and regulation by the *εLCY* 5’UTR RNA structural switch. β-branch carotenoid levels correlate with *εLCY* expression levels. Metabolic feedback triggered during etioplast, and chloroplast development regulates *εLCY* expression in order to balance α/β-carotene homeostasis during seedling morphogenesis and leaf development. β-branch carotenoids are reduced by the loss-of-function CRTISO mutant (*ccr2*) and by norflurazon (NFZ; inhibits PHYTOENE DESATURASE, PDS) treatment of seedlings. Metabolic feedback via a retrograde signal (e.g., heme or an apocarotenoid signal; ACS) generated in the plastid signals the nucleus to regulate *εLCY* levels. Mutations that stabilise an alternative distinct RNA structural probability (shape-1) of the *εLCY* 5′UTR can establish a post-transcriptional expression platform that reduces protein activity without affecting transcript levels. The shape-1 5′UTR fragment harbours an aptamer domain capable of sensing a feedback signal generated in *ccr2* or in NFZ-treated seedling tissues that can repress reporter gene expression. The *εLCY* 5′UTR represents a new conformational RNA regulatory switch that could modulate carotenoid biosynthesis through the branch in the pathway to maintain metabolic homeostasis during plastid biogenesis.

### The *εLCY* promoter modulates feedback regulation during carotenogenesis

We investigated the mechanism by which metabolic feedback regulates *εLCY* expression. Retrograde signalling molecules can alter transcription and/or post-transcriptional processes to moderate metabolic feedback regulation in various plastid types (Hernandez-Verdeja and Strand, 2018; Moreno et al., 2021; Wu and Bock, 2021). Our analysis of two-independent isogenic WT and *ccr2* transgenic lines harbouring a -450 bp *εLCY* promoter fragment fused to the *FiLUC* reporter gene revealed that reporter levels mirrored *εLCY* expression during genetic, environmental, and chemically induced metabolic feedback regulation. That is, transcription enabled by the *εLCY* promoter was suppressed by ∼50% in *ccr2* compared to WT during skoto-morphogenesis and substantially enhanced by 4.5-fold during photomorphogenesis. Similarly, when NFZ was used to block β-branch carotenoid biosynthesis in both dark- and light-grown WT seedlings, the *εLCY*::*FiLUC* activity declined. Seedling de-etiolation and NFZ treatment did not alter CaMV35s::*FiLUC* expression or activity. Therefore, the -450bp *εLCY* promoter fragment conferred similar changes in reporter gene expression and activity in response to metabolic feedback triggered in *ccr2* and NFZ-treated WT seedlings during both skoto- and photo-morphogenesis.

### The *εLCY* 5’UTR has an IRES structural motif and posttranscriptional expression platform

We interrogated the *εLCY* -450 bp promoter searching for features that could modulate metabolic feedback regulation. Our 5′RACE analysis identified three distinct TSSs within the first -205 bp (TSS_1_) upstream from the *εLCY* start codon that separated three conserved domains similarly positioned relative to the *εLCY* translation initiation codon from other *Brassicaceae* species. Inr elements contained within CD-1 and adjacent CCAAT box located slightly upstream of TSS_1_, could associate with three pyrimidine patch (YP) motifs downstream of each TSS to alternate transcription between the three fragments. The longest TSS_1_ (-205 to -200 bp) and shortest TSS_3_ (-75 to -70 bp) 5′UTR fragments were barely detected in etiolated seedlings, while TSS_3_ was prominent in light-grown seedlings. In contrast, TSS_2_ (-143 to -130 bp) had adjacent Inr (-152 to -147 bp) and overlapping IME-Like (-130 to 137 bp) elements that could recruit RNA polymerase II to enrich this fragment in both dark and light-grown seedlings (Parra et al., 2011; Laxa et al., 2016). However, gain-of-function and deletion studies indicated that *cis*-acting motifs upstream of the shorter 5′RACE validated TSS_3_ fragment or present in the longer/abundant TSS_2_ fragment do not impact CaMV35s enabled FiLUC activity levels in photosynthetic leaf tissues from transgenic Arabidopsis and agrobacterium-infiltrated tobacco plants.

We identified a conserved IRES structural motif (-82 bp) shown to utilise a cap-independent mechanism in promoting translation (Martinez-Salas et al., 2018). IRES motifs are evolutionarily conserved among viral mRNAs and found in eukaryotes where they impact upon RNA structure to recruit ribosomes, initiation factors and/or RNA-binding proteins (RBPs). Removing 3′ or 5′ segments of the *εLCY* IRES (-82 bp) did not affect CaMV35s-enabled FiLUC activity in photosynthetic leaf tissues. This was not surprising as IRES RNA molecules need to acquire a structural conformational change to impart translation activity and regulate responses to chemical ligands and/or environmental signals (Lozano et al., 2015; Martinez-Salas et al., 2018).

Viral IRES elements can be conformationally flexible RNA switches, whose state can be captured by the binding of a common ligand and are smaller than previously discovered RNA riboswitches (Boerneke et al., 2014). The nine mutations within the *εLCY* 5′UTR Shape-1 fragment stabilised a single RNA structural definition surrounding the IRES and repressed FiLUC activity rather than transcript levels. Shape-7 complemented seven of the nine mutations leaving two targeting the stem of hairpin-1/hairpin-2 within CD-3 that repressed FiLUC activity without altering transcript levels enabled by the CaMV35s promoter. RBPs have the capacity to assemble ribonucleoprotein complexes on RNA elements such as the IRES from the time of synthesis to degradation and modulate pre-mRNA processing, transcript stability, post-transcriptional and translational regulation (Fitzgerald and Semler, 2009; Martínez-Salas et al., 2013; Yang and Wang, 2019).

The post-transcriptional repression caused by shape-1 could arise from the formation of a novel *cis*-acting element or structural motif capable of recruiting a *trans*-acting RBP. However, deleting the sequence motif harbouring hairpin-1 did not impact FiLUC activity. The shape fragments showed negligible separation between the structure prediction curves in the mountain plot representations. All the shape mutations impaired formation of hairpin-1 and hairpin-2, while stabilising the probability of hairpin-3 in 85% of shape-7 structural definitions. It appears most likely that a structural definition change, rather than the introduction of a mutation or novel *cis*-acting motif in the *εLCY* 5′UTR modulates FiLUC activity levels. The stabilisation of hairpin-3 could resemble a terminator-like structure associated with the IRES structure that impedes the recruitment of RBPs required to promote efficient translation. The *εLCY* 5′UTR expression platform might be capable of switching between conformational RNA structural definitions. One that maintains permissive translation, and an alternative structural definition that establishes a terminator-like feature within CD-3 upstream of the IRES expression platform to impede translation (Figure 7).

### The *εLCY* 5’UTR harbours an aptameric domain responsive to metabolic feedback signalling

We examined if the *εLCY* 5′UTR region within the *εLCY* promoter could mediate feedback regulation. Riboswitches are commonly found in bacteria and a single TPP-riboswitch has been conserved in plants whereby it can sense thiamine levels and affect RNA 3′ end processing to control gene expression (Wachter et al., 2007; Etzel and Mörl, 2017). A typical RNA riboswitch has an aptamer domain that binds the ligand, and an expression platform that undergoes structural changes to regulate either initiation, elongation, or termination of transcription and/or translation (Assmann et al., 2023). IRES RNA motifs can also be functionally conserved switches involved in viral IRES-driven translation and constitute a new paradigm for ligand captured switches that differ from metabolite-sensing riboswitches with regard to their small size, intrinsic stability, and structural definition of the constitutive conformational states (Boerneke et al., 2014). Unlike transgenic lines harbouring the *εLCY* promoter-reporter fusion, WT seedlings harbouring the CaMV35s-5′UTR::FiLUC reporter were not responsive to NFZ mediated metabolic feedback inhibition. The most probable 5′UTR structural probability (69%) appears to not display a structural definition of a metabolite-binding aptameric domain. In contrast, the shape-1 fragment that had a stabilised structural definition and terminator-like structure within CD-3 has the integrity of a metabolite-binding aptameric domain since NFZ treatment suppressed *FiLUC* transcript and activity levels in WT de-etiolated seedlings. *ccr2* de-etiolated seedlings harbouring the CaMV35s-shape-1::*FiLUC* transgene similarly displayed lower reporter activity levels in comparison to WT lines, which can be attributed to the loss-of-function in CRTISO that signals metabolic feedback (Cazzonelli et al., 2020). How the presence (or absence) of a signal, whether derived from a β-branch carotenoid substrate or not, and how it interacts with the aptamer domain of shape-1 to modulate the expression platform has yet to be established. The shape-1 structural definition of the *εLCY* 5’UTR should impart an aptameric domain capable of sensing metabolic feedback during plastid biogenesis to regulate reporter gene expression (Figure 7).

### The *εLCY* 5’UTR resembles a metabolite-responsive RNA regulatory switch

The *εLCY* 5′UTR appears to impart a conformational RNA regulatory switch having sequence homology with other *Brassicaceae* species, a viral IRES structural motif proximal to the start codon and exhibits structural definitions with switching probabilities similar to that of ancient bacterial metabolite-binding riboswitches and viral IRES ligand-responsive RNA switches. By modelling 27 different classes of experimentally validated metabolite-binding riboswitches, we found the *εLCY* 5′UTR can form two distinct structural definitions representing 69% and 29% of probability distributions represented by dot-bracket notations keeping consistent with *in silico* analysis demonstrating that 42% of riboswitches displayed a minimum of 20% to maximum of 78% probability for a representative structural configuration. Intriguingly, the *εLCY* IRES structural motif shared weak sequence homology with the SMK_box metabolite-binding RNA riboswitch. Conformational switching plays a role in the function of the IRES elements (Boerneke et al., 2014). RNA structural bioinformatics identified 1) an optimal negative mfe ranging from -21 to -37 kcal/mol, 2) a centroid structural peak with lower minimal entropy located within the putative IRES/SMK_Box region that harbours fewer secondary structure outcomes within this space, 3) a lower fit of greater separation between the structure prediction curves (centroid, partition function; pf, and mfe) surrounding three distinct RNA stem-loop terminator-like structures within CD-3, and 4) distinct and separate energy clouds by morphological distance plot, string distance, and clustering validation reminiscent of a new RNA structural switch in plants.

The alternative shape-1 structural variant of the *εLCY* 5’UTR may modulate two functions to, 1) sense metabolic feedback and represses *FiLUC* transcript levels probably via an aptamer domain within its structural definition, and 2) regulate protein activity levels indicative of a post-transcriptional expression platform within its structural definition. The transcriptional and post-transcriptional regulations displayed by the aptamer and expression domains respectively, are not exclusive and indicate a role for RBP in modulating metabolic feedback. The *εLCY* 5’UTR could represent a conformational RNA structural switch that has evolved to fine-tune flux through the branch in carotenoid biosynthesis and maintain metabolic homeostasis during plastid biogenesis (Figure 7).

## MATERIALS & METHODS

### Plant germplasm

All Arabidopsis germplasm are in the Columbia (Col-0) ecotype. Germplasm previously generated by other studies include: *ccr2.1*/*crtiso* (Park et al., 2002), ziso#11C (*zic1-3*: Salk_136385) (Chen et al., 2010), *det1-1* (CS6158) (Chory et al., 1989), *det1-154* (Cazzonelli et al., 2020), *ccr2 det1-154* (Cazzonelli et al., 2020), *ccd4* (Salk_097984c), *ccr2 ccd4* (Cazzonelli et al., 2020), and MicroTom *tangerine^Mic^* (Kachanovsky et al., 2012).

### Plant growth media, seed sterilisation and chemical treatment assays

Arabidopsis seeds were stratified at 4-5°C for 2-3 days to synchronise germination. Soil-grown Arabidopsis plants were sowed on DEBCO seed raising mix with Osmocote fertilizer (30 mL/ 5L) within growth cabinets (Climatron, Thermoline Scientific) or walk-in growth rooms fitted with fluorescent lights (120-150 μE m^−2^ s^−1^) and maintained under 16/8 hr day/night photoperiod at 21°C constant temperature. For artificial media-grown seedlings, seeds were first surface sterilised for 3-4 hours by chlorine gas treatment (produced from mixing 100 mL 4% sodium hypochlorite and 3 mL of 37% w/v HCl), rinsed in 70% ethanol, followed by sterile H_2_O. Arabidopsis and tomato etiolated seeds were sown within a petri dish plate containing MS media (Caisson, MSPO1) supplemented with 0.5x Gamborg’s vitamin solution (0.5 mL/L) (Sigma-Aldrich), and 0.5% phytagel (Sigma-Aldrich) (5 g/L). Plates were wrapped in a double-layered aluminium foil and incubated in a dark growth cabinet at 21°C and 60% constant humidity for 7-days after stratification. For de-etiolation experiments, Arabidopsis seeds were stratified and grown in the dark at 21°C for 4 days prior to exposed seedlings to constant low fluorescent lighting (80-90 μE m^−2^ s^−1^) for 72 hr at 21°C, unless otherwise indicated. Etiolated tissues were harvested under a non-photosynthetic dim green LED light in a dark room. NFZ treatment of seedlings was performed by growing seedlings in MS media supplemented with 5 μM NFZ (dissolved in 100% DMSO).

### Molecular vector construction

pT*εLCY*Prom-*εLCY*5′UTR::FiLUC (previously referred to as pT*eLCY*-4F::FiLUC) harbours a - 450bp region (promoter plus 5’UTR) cloned upstream of the *FiLUC* gene into pTm35:FiLUC (digested with *Xba*I/*Nco*I to remove the minimal 35s promoter), as previously described (Cazzonelli et al., 2010). The -133 bp *εLCY* 5’UTR and series of *εLCY* 5’UTR mutated and deleted (MutDels) sequence variants downstream of a min35s domain that incorporated 5’ *Xho*I and 3’ *Nco*I restriction sites were synthesised and cloned into the pMat intermediate plasmid (Life Technologies, Germany). Fragments were subsequently removed from pMat and cloned into pT35enh::*FiLUC* binary vector using *Nco*I/*Xho*I restriction endonuclease digestion and T4 ligation as previously described (Cazzonelli and Velten, 2006). This generated nine MutDels constructs where the *εLCY* 5’UTR was fused downstream of the CaMV35 promoter and upstream of the luciferase reporter gene (Figure 5D).

The pT*35S*-*εLCY*5′UTR-Shape1::FiLUC vector was further modified by sequentially restoring mutated sequences to generate three new constructs referred to as "Mut2Y" (Shape-2), "Mut6Y" (Shape-6) and "Mut7Y" (Shape-7). The subsequent new destination vector was modified such that *Hind*III sites flanking the 35Senh and min35s domain were replaced to introduce an internal *XbaI* site after the min35s domain and internal to the *Hind*III site. In brief, the region between 35Smin and *εLCY* 5′UTR insert was amplified using Platinum® Pfx DNA polymerase (Invitrogen) with HindIII_T35_F1 and HindIII+XbaI_T35_R1 primers and the amplicon cloned into a pCR4-TOPO intermediate vector. pCR4-TOPO::35Smin::*εLCY*5′UTR was digested with *HindIII,* and the purified T35Smin-*εLCY*5’UTR fragment ligated into the pT35s-*εLCY*5′UTR::FiLUC vector digested with *HindIII* and treated with FastAP Thermosensitive Alkaline Phosphatase (1 U/μL) (ThermoFisher) to create pT35sX- *εLCY*5′UTR::FiLUC. The new MutDels of the *εLCY*5′UTR fragments were amplified using 1F_Mut5UTR, 4F Mut5UTR, 6F_Mut5UTR plus 1R_Mut5UTR primers (harbour flanking 5’ *XbaI* and 3’ *NcoI* sites) and pT35s-*εLCY*5′UTR::*FiLUC* as DNA template to generate amplicons for Mut2Y, Mut6Y and Mut7Y, respectively. All amplicons were cloned into pCR^®^4-TOPO^®^ and validated by Sanger sequencing. Mut2Y, Mut6Y and Mut7Y amplicons were subsequently digested from the intermediate vector and ligated into the modified pT35sX-*εLCY*5′UTR::FiLUC vector to create pT35S-εLCY5′UTR-Mut2Y::FiLUC, pT35S::εLCY5′UTR-Mut6Y::FiLUC, and pTV35S-εLCY5′UTR-Mut7Y::FiLUC binary constructs using an *Xba*I/*Nco*I cloning strategy.

The bacterial CrtI gene from *Pantoea agglomerans* (*PaCrtI*) was fused downstream of the Arabidopsis small subunit (SSU) of RuBisCo transit peptide (56 amino acids) (Wong et al., 1992) controlled by the CaMV35S enhancer to create the binary vector pT35S::*PaCrtI*. The nopaline synthase terminator (NosT) within pT35enh::*FiLUC* (Cazzonelli et al., 2010) was re-amplified to include additional 5’ (*Bgl*II and *Avr*II) and 3’ (*Pst*I) restriction sites, cloned into the TOPO-Blunt-II intermediate vector (Invitrogen), and subsequently used to replace the original NosT to create pT35enh::*FiLUC-*NosTMod. The SSU-*PaCrtI* gene was chemically synthesized and incorporated bordering ’5’ (*Nco*I) and 3’ (*Avr*II) restriction sites cloned into the pMat intermediate plasmid (Life Technologies, Germany). The *FiLUC* gene was excised from pT35enh::*FiLUC-*NosTMod using *Nco*I/*Avr*II and replaced with *PaCrtI* to create the pT35enh::SSU-*PaCrtI* binary vector.

### Plant Transformation and selection of transgenic lines

Arabidopsis transformation with Agrobacterium harbouring binary plasmids was performed using a modified floral dip method (Cazzonelli and Velten, 2008). Agrobacterium and the containing binary plasmid were selected on Rifampicin (25 μg/L) and spectinomycin (100 μg/L). The TDNA within the binary vectors harboured *PHOSPHINOTHRICIN ACETYL TRANSFERASE* (*BAR*) that allowed the selection of transgenic and homozygous lines by spraying plants with BASTA herbicide (50 mg/L) as previously described (Cazzonelli and Velten, 2004). The generation of pTm35enh::*FiLUC* transgenic WT lines was previously described (Cazzonelli et al., 2010). For exogenous expression of *PaCrtI* in *ccr2*, nineteen independent transgenic lines haboring T35enh::*SSU-PaCrtI* survived BASTA herbicide selection. These lines were grown on MS media containing 2.5μM NFZ and classified as green, pale-green, or albino-green (i.e. sectored) in phenotype, while non-transformed WT plants were completely albino in appearance. A homozygous line (T35enh::*SSU-PaCrtI*#11) that displayed a green phenotype and segregated in a typical 3:1 mendelian manner (Chi-squared test) was chosen to be the representative line.

### *E. coli* transformation, electroporation and agro-infiltration

Chemically competent *E. coli* (DH5α) cells were prepared as previously described (Holsters et al., 1978). Plasmids were transferred into competent *E. coli* cells using a conventional heat-shock (42 °C) transformation method. Electrocompetent Agrobacterium (LBA4404) cells were prepared as previously described (Hanahan, 1985). Plasmids were electroporated into electro-competent Agrobacterium using a Bio-Rad MicroPulser (Bio-Rad, Hercules, USA) according to the manufacturer’s protocol. Agro-inoculation was performed using infiltration media (10 mM MgCl_2_, 100 μM acetosyringone and 1 mM MES (2-(N morpholino) ethanesulfonic acid) at pH 5.6 via needless syringe as previously described (Cazzonelli and Velten, 2004).

### Luciferase bioassays

Luciferase assays were performed as described (Velten et al., 2008), and used to select representative transgenic lines as well as distinguish between homozygous and heterozygous lines based upon luminescence (*FiLUC* activity) emitted from identical-sized leaf disks collected from mature leaves. Individual leaf disks were placed on 35μL assay media (50 mM MES (pH 5.6), 0.5% glucose, 2 mM NaPO_4_, 0.5% v/v DMSO) in individual wells of a 96-well plate (Corning^®^ NBS). Then 15 μL of luciferin assay media (1 mM luciferin, 50 mM MES (pH 5.6), 0.5% glucose, 2 mM NaPO_4_, 0.5% v/v DMSO) was added prior to quantifying luminescence as previously described (Cazzonelli and Velten, 2006) using the CLARIOstar^®^ *Plus* plate reader (BMG LABTECH). Luminescence imaging of whole plants was performed by incubating them in 0.5 mM luciferin for 10 minutes prior to imaging light emission using a CCD camera (Model DV435; Andor Technologies) and ImagePro Plus 4.5.1 software (Media Cybernatics). Captured images of luminescence were enhanced and overlayed using NIH Image J software (version 1.53).

### Total RNA isolation and cDNA synthesis

Frozen leaf samples were physically disrupted using TissueLyser (Qiagen). The total RNA was extracted from the fine powder of plant tissue with Spectrum™ Plant Total RNA kit (Sigma-Aldrich) according to the manufacturer’s protocol. Isolated total RNA samples were quantified with NanoDrop 2000 and used for cDNA synthesis after validating their integrity on a 1% TBE agarose gel. The total RNA (1 to 2 μg) was used to synthesise cDNA with the Bioline Tetro cDNA synthesis kit by following the manufacturer’s instructions. The reaction mix was prepared with 1 μL Oligo (dT), 1 μL 10mM dNTP mix, 4 μL 5x RT Buffer, 1 μL RiboSafe RNase Inhibitor, 1 μL Tetro Reverse Transcriptase (200 U/μL), 1-2 μg total RNA and DEPC-treated dH_2_O up to 20 μL total reaction volume. The mixture was incubated at 45°C for 30 min, then at 85°C for 5 min to terminate and chilled on ice for 5 min.

### Quantification of relative gene expression and PCR

The relative transcript abundance in plant tissues was quantified using LightCycler 480 (Roche, Australia). Primers were designed according to the sequence information of organisms in the NCBI GenBank and TAIR databases using Primer3Plus (Untergasser et al., 2012). Quantitative reverse transcription PCR (qRT-PCR) was performed with the mixture of 2 μL of primer mix (2 μM from each F & R primer), 1 μL 1/10 diluted cDNA template, 5 μL LightCycler 480 SYBR Green I Master mix and distilled water up to the total volume of 10 μL. For each sample, three technical replicates for each of one to three biological replicates were tested. The relative gene expression levels were calculated by using relative quantification (Target Eff ^Ct(Wt-target)^ /Reference Eff ^Ct(Wt-target)^) and fit point analysis (Pfaffl, 2001). The *Protein Phosphatase 2A* (*PP2A*, At1g13320) and/or *TIP41* (At4g34270) were used as house-keeper reference control genes for all qRT-PCR experiments (Czechowski et al., 2005). All primer sequences are listed in Table S3.

General PCR reactions were performed by preparing a mixture with 2.5 μL 10 X Taq polymerase buffer, 1 μL dNTP (10 mM), 2 μL MgCl_2_ (10 mM), 0.25 μl Taq DNA polymerase (5 U/μL), 1 μL/each of forward (F) and reverse (R) primers (10 μM/μL), 1 μL template DNA (∼100 ng/μL) and dH_2_O up to 25 μL. Thermal-cycler adjustments were: 5 min 95°C pre-denaturation followed by 35 cycles of 1 min at 95°C, 1 min at 45-65°C 1 min at 72°C and 5 min for the final extension at 72°C. The annealing temperature was variable depending on the used primers. 2% TBE agarose gel was used to confirm the length of each reaction product.

### 5′ Rapid Amplification of cDNA Ends (5’RACE)

The integrity of total RNA samples used for Rapid Amplification of cDNA Ends (RACE) assays was validated using Agilent 2100 Bioanalyzer in WSU’s NGN facility. 5′RACE assays were performed using the GeneRACER kit according to the manufacturer’s protocol. RACE cDNA libraries were prepared with SuperScript III Reverse Transcriptase using GeneRacer™ Oligo dT Primer according to the manufacturer’s protocol. 5′ ends of gene-specific cDNAs were amplified with Platinum^®^ Pfx DNA polymerase (Invitrogen) and subsequently cloned into pCR^®^4-TOPO^®^ for Sanger sequencing. The sequences of gene-specific primers used in the first step of amplification and the nested PCR can be found in Table S3.

### Carotenoid Purification and HPLC Analysis

Frozen tissues were homogenised to a fine powder using TissueLyser® (QIAGEN), while root tissues, on the other hand, were homogenised using mortar and pestle. Carotenoids were extracted as previously described (Alagoz et al., 2020) using a reverse-phase high-performance liquid chromatography (HPLC) (Agilent 1260 Infinity) and a YMC-C30 (250 × 4.6 mm, S-5µm) column. Carotenoids and chlorophylls were identified based on their retention time relative to known standards and their light emission absorbance spectra at 440 nm (chlorophyll, β-carotene, xanthophylls, pro-neurosporene, tetra-*cis*-lycopene, other neurosporene and lycopene isomers), 400 nm (ζ-carotenes), 340 nm (phytofluene) and 286 nm (phytoene). Absolute quantification of carotenoid and chlorophyll pigments was performed as described (Anwar et al., 2022).

### *In-silico* bioinformatics analysis

*In-silico* prediction of RNA secondary structure formations was performed using desktop and online versions of RNAShapes (https://bibiserv.cebitec.uni-bielefeld.de/rnashapes) (Steffen et al., 2006; Janssen and Giegerich, 2015) and RNAfold (Vienna package; http://rna.tbi.univie.ac.at/cgi-bin/RNAWebSuite/RNAfold.cgi) (Grillo et al., 2010; Lorenz et al., 2011). UTRscan (http://bioinformatics.ba.itb.cnr.it/?Software__UTRscan) and IRESite (https://iresite.org) were used to assess sequence homology against a UTR database and predict functional elements and validate against a database of experimentally verified IRES (Pesole and Liuni, 1999; Mokrejs et al., 2010). cis-acting regulatory elements/motifs were identified in the - 450 bp *εLCY* promoter region using PlantCare (Lescot et al., 2002). NCBI Blast (https://blast.ncbi.nlm.nih.gov/Blast.cgi) and Primer3Plus (version 2.4.0) (https://primer3plus.com) (Untergasser et al., 2012) were used to identify gene sequences and to design qPCR primers, respectively. Geneious (version 10.2.6) was used to perform the sequence alignments. CLC Sequence Viewer (version 8.0.0) *in-silico* cloning editor was used to design the plasmids. Genevestigator was used to interrogate the effect of NFZ on gene expression in WT seedlings grown under light (Hruz et al., 2008). Arabidopsis eFP browser was used to interrogate the differential expression of *εLCY* in young and mature leaves, using data from a previous study (Winter et al., 2007; Klepikova et al., 2016). Statistical analysis for qPCR and HPLC results were performed with SigmaPlot (v14.0) using one-way or two-way ANOVA with post hoc Tukey (HSD), depending on the experimental setup. Figures were designed using a commercial software Adobe Illustrator.

## Abbreviations

ACS: apocarotenoid signal
β-ACS: β-carotene derived apocarotenoid signal
*cis*-ACS: *cis*-carotene derived apocarotenoid signal
FiLUC: intron modified *FIREFLY* (*Photinus pyralis*) *LUCIFERASE* gene
MutDels: mutations and/or deletions
NFZ: norflurazon
Nos^t^: nopaline synthase terminator
uORF: upstream open reading frame
RLU: relative light units
TSS: transcription start site
5’UTR: 5’ Untranslated Region

## SUPPLEMENTARY FIGURE LEGENDS

**Figure S1. Absolute levels of linear *cis*-carotenes in *ziso*, *ccr2 ziso*, and *ccr2* etiolated tissues exogenously expressing *PaCrtI*.** (**A**) Linear *cis*-carotene levels in *ccr2*, *ziso*, and *ccr2 ziso* etiolated seedlings. (**B)** Linear *cis*-carotenes and (**C**) cyclic carotenoids in etiolated tissues of *ccr2* and *ccr2* T35enh::SSU-*PaCrtI#11*. Significant differences are denoted by lettering (Two-way ANOVA) or stars (one-way ANOVA; *p <0.05, **p <0.005, ***p <0.001). Data are representative of three separate experiments, and bars define standard error (n=3-4). Abbreviations: violaxanthin (Vio), neoxanthin (Neo), antheraxanthin (Ant), lutein (Lut), β-carotene (β-car), pro-neurosporene (P-Neu), prolycopene (P-Lyc), 9,15,9’-tri-*cis*-ζ-carotene (tc-ζc), 9,9’-di-*cis*-ζ-carotene (dc-ζc), 15-*cis*-phytofluene (Phf), 15-*cis*-phytoene (Phy).

**Figure S2. Loss-of-function in CCD4 enhances β-carotenoid accumulation in WT and *ccr2* etiolated seedlings.** Quantification of carotenoids in dark-grown seedlings. **A**) Cyclic carotenoid levels in WT and *ccd4* etiolated cotyledons. **B**) Linear *cis*-carotene levels in WT and *ccr2 ccd4* etiolated cotyledons. Significant differences are denoted by stars (ANOVA; *p <0.05, **p <0.001). Data represents multiple experiments, and bars denote standard error (n=3-4). Abbreviations: Violaxanthin (Vio), Neoxanthin (Neo), Antheraxanthin (Ant), Zeaxanthin (Zea), Lutein (Lut), β-carotene (β-car), pro-neurosporene (P-Neu), prolycopene (P-Lyc), 9,15,9’-tri-*cis*-ζ-carotene (tc-ζc), 9,9’-di-*cis*-ζ-carotene (dc-ζc), 15-*cis*-phytofluene (Phf), 15-*cis*-phytoene (Phy).

**Figure S3. Carotenoid content in norflurazon-treated wild-type and *ccr2* etiolated seedlings**. **A-B)** HPLC chromatogram (A) showing carotenoid traces (A_440nm_) and absolute levels (B) of phytoene (A_286nm_) as well as phytofluene (A_348nm_) from WT and *ccr2* dark-grown seedlings grown on artificial media with (+) or without norflurazon (NFZ; 5 µM). Significant differences are denoted by lettering (Two-way ANOVA). **B)** Representative image of a norflurazon (NFZ; 5 µM) treated and untreated (control) wild type (WT) seedling. Data represent three separate experiments, and standard error bars are displayed (n=3-4). Abbreviations:, violaxanthin (V), neoxanthin (N), antheraxanthin (A), lutein (L), zeaxanthin (Z), β-carotene (βc), pro-neurosporene (P-Neu), prolycopene (P-Lyc), 9,15,9’-tri-*cis*-ζ-carotene (tc-ζcar), 9,9’-di-*cis*-ζ-carotene (dc-ζcar),.

**Figure S4. Pigment content and *εLCY* expression in young and old leaves**. **A)** Carotenoid levels in WT and *ccr2* de-etiolated seedlings following 72 hrs of constant illumination. **B)** Absolute expression *εLCY* levels in young and older mature leaf tissues as depicted by the TAIR eFP browser (Klepikova Atlas). **C**) Carotenoid and chlorophyll levels in old mature and young emerging leaves from the 3-weeks-old rosette leaves. Data are representative of three separate experiments, and bars define standard error of the mean (n=3-4). Significant differences are denoted by lettering (Two-way ANOVA) or stars (one-way ANOVA; *p <0.05, **p <0.005, ***p <0.001). Abbreviations: violaxanthin (Vio), neoxanthin (Neo), antheraxanthin (Ant), lutein (Lut), β-carotene (β-car), chlorophyll a (Chla), chlorophyll b (Chlb).

**Figure S5. Luciferase activity and transcript levels in transgenic plants harbouring the *εLCY* and CaMV35s promoter-reporter gene fusions.** A-B) Independent transgenic lines harbouring *εLCY*Prom-*εLCY*5′UTR::FiLUC (A) and *CaMV35S*::FiLUC (35S) (B) transgene fusions. Luciferase (FiLUC) activity was determined by luminescence quantified as relative light units; RLU) from equal-sized leaf discs (two leaf discs per leaf) dissected from independent transgenic lines (T_1_ plants) using the *in vivo* floating leaf-disk assay (Cazzonelli and Velten, 2006). Independent lines are ordered by increasing luciferase activity along the x-axis and representative lines from the *εLCY* (εLP#2e; εLP#4c) (A) and *CaMV35S* (35s#B30, 35s#C1) (B) promoter lines are displayed. (**C**) Relative expression of *εLCY* and *FiLUC* transcript levels in transgenic 35s#C1 etiolated and de-etiolated seedling tissues. Significant differences are denoted by lettering (ANOVA).

**Figure S6. Deletions, mutations, *cis*-acting motifs and conserved domains in the *εLCY* promoter and 5′UTR. A)** *cis*-acting regulatory elements identified in the -450 bp *εLCY* promoter region using PlantCare (Lescot et al., 2002), UTRScan and IRESite (Pesole and Liuni, 1999; Mokrejs et al., 2010) bioinformatics software. The putative elements, Downstream Promoter Element (DPE), Intron-Mediated Enhancer (IME), Initiator (Inr), and Internal Ribosome Entry Site (IRES) elements were identified by aligning the *εLCY* promoter sequence using Clustal Omega alignment software. Transcription Start Sites (TSS) and conserved domains are marked upstream of the start codon (ATG). An upstream open reading frame (uORF) consisting of 2 amino acids (Met, Val) is denoted by a red underline (-47 to -39 bp). Nucleotide positions shown refer to the length of *εLCY* 5′UTR fragments. **B)** Sequence alignment of *εLCY* 5′UTR mutation and deletion (MutDels) fragment variants. Alignment was performed using Geneious Prime Software (v10.2.6). The sequence logo shows the mutations introduced. The font size reflects the abundance of the nucleotide. Nucleotide positions indicated above the sequence logo reveal the position of relevant MutDels. The red vertical box signifies the two mutated nucleotides in shape fragments that alter 5′UTR RNA structural definitions and CaMV35S promoter enabled reporter activities.

**Figure S7. Representative RNA secondary structural plots and prediction probabilities**. Vienna RNA structural representations generated by RNAShapes using dot-bracket notation for the *εLCY* 5′UTR (A), shape-1 (B), shape-2 (C), shape-6 (D), and shape-7 (E) sequences. Red lines and circled DNA base pairs denote the hairpin-1, hairpin-2 and hairpin-3 structural definitions in the *εLCY* 5′UTR (A). The percentage (%) value displayed above structural definitions reflects the highest percentage shape probability differentiating the dot-bracket notations by hairpin definitions with CD-3 (Figure S7).

**Figure S8. *In silico* RNA structural analysis of the *εLCY* shape variants modulating a post-transcriptional expression platform.** (**A**) RNAfold mountain plots showing RNA secondary structure predictions for *εLCY* 5’UTR, shape-1, shape-2, shape-6, and shape-7. Plots represent the mfe structure, thermodynamic ensemble of RNA structures (pf), and the centroid structure in a plot of height m(k) versus position (bp), where the height is given by the number of base pairs enclosing the base at a position. "mfe" represents minimum free energy structure; "pf" indicates partition function; "centroid" represents the best average structure. Loops correspond to plateaus (hairpin loops are peaks), and helices to slopes. RNAfold calculates the entropy of each bp along the RNA sequence, indicating the stability of the RNA secondary structure per given sequence (starting at -1 to -133 bp) which is shown below the mountain plot. Low entropy values indicate high stability of RNA secondary structures indicated by the mountain height/bp. **(B)** RNAShapes analysis of the 5′UTR, shape-1, shape-2, shape-6, and shape-7 sequences showing Vienna shape representatives (dot-bracket notation), percentage shape probability (Prob.) and minimal free-energy (MFE; kcal/mol) associated with secondary structure formation (Steffen et al., 2006; Janssen and Giegerich, 2015). Dot-bracket notation for RNA secondary structures is represented by a string length of matching brackets and dots, where a base pairing is represented by a ’(’ and unpaired bases are represented by dots ’.’ (Gruber et al., 2008). Green or red dot-brackets denote similar structural representations. The structural probability (prob.) >0.01 (1%) and minimum free energy (MFE) are displayed. Red and orange lines above the *A. thaliana* 5’UTR sequence denote CD-3 and IRES motif, respectively. The boxed regions denote three stem-loop hairpin structures found in Conserved Domain-3 (CD-3). (**C**) Base-pairing probability and (**D**) positional entropy of the most probable structures for *εLCY* 5’UTR, shape-1, shape-2, shape-6, and shape-7 as generated by RNAfold software using default settings. A base pairing probability value of 1 means it is highly stable. A positional entropy value of 0 indicates the base-pair is always unpaired or paired with its partner.

## SUPPLEMENTARY TABLE LEGENDS

**Table S1.** Correlations between the regulation of *εLCY* and carotenoid levels in plants.

**Table S2.** Carotenoid percent composition in rosette leaf tissues from *ccr2* lines transformed with 35Senh::*SSU-PaCrtI*

**Table S3.** Primers used for qPCR, cloning, and 5’Rapid Amplification of cDNA Ends.

## SUPPLEMENTARY DATA FILE LEGENDS

**Data File S1.** Nucleic acid sequence-based homology analysis of *εLCY* 5′UTR for screening similarity with Rfam riboswitch database.

**Data File S2.** The shape-switching potential of 27 different riboswitch classes found in the Rfam database. Analysis was performed based on RNA structures with minimum free energy and maximum stability. The number of shapes with >10% minimum probability is considered significant.

## FUNDING

This work was financially supported by the Australian Research Council Discovery Grant DP130102593 awarded to CIC and BJP.

## ACKNOWLEDGEMENTS

YA and JJN were supported by a Western Sydney University-Hawkesbury Institute for the Environment Postgraduate Scholarship Award. We acknowledge Prof. Ralph Bock and Dr Christian Schudoma from the Max-Planck Institute for assisting with the RNA infernal-scan Rfam covariance model analysis of the 5’UTR.

## AUTHOR CONTRIBUTIONS

CIC conceived ideas and YA and JJN designed research. YA, JJN, JW, SH, and CIC performed experiments, analysed data, and prepared figures/tables. RA, YA, and CIC performed the RNA structural and riboswitch bioinformatics analysis. YA and CIC wrote the manuscript. Detailed contributions are as follows; Figure 1 (YA, JJN, JW, CIC), Figure 2 (YA, JJN, RA, CIC), Figure 3 (YA, CIC), Figure 4 (YA, JJN, SH, CIC), Figure 5 (JJN), Figure 6 (JJN), Figure 7 (YA, JJN, CIC), Figure S1 (YA, JJN), Figure S2 (YA), Figure S3 (YA), Figure S4 (YA), Figure S5 (YA, CIC), Figure S6 (YA, JJN, CIC), Figure S7 (CIC), Figure S8 (CIC), Table 1 (CIC, RA), Table S1 (YA), Table S2 (JA, CIC), Table S3 (YA), Data File S1 (CIC), Data File S2 (RA, CIC). CIC was the primary supervisor of YA, JJN, JW and SH. BJP co-supervised JW and SH. DT co-supervised YA. All authors have read and approved the manuscript.

## Notes

The authors declare no conflict of interest.

